# Presynaptic temporal dynamics flexibly set input weights in the mouse escape circuit

**DOI:** 10.64898/2026.05.18.724906

**Authors:** Yu Lin Tan, Ashviniy Thamilmaran, Natalia Zernicka-Glover, Dario Campagner, Tiago Branco

**Affiliations:** UCL Sainsbury Wellcome Centre for Neural Circuits and Behaviour, London, United Kingdom; UCL Gatsby Computational Neuroscience Unit, London, United Kingdom

## Abstract

Animals facing threat must integrate multiple streams of information — about danger, environment, and internal state — into a time-pressured escape decision. In mice, this computation is performed by glutamatergic neurons of the dorsal periaqueductal grey (dPAG), but how their convergent long-range inputs combine to drive flexible decisions is unknown. Here we find that the functional weight of each input is set predominantly by the temporal statistics of its presynaptic activity, rather than by pathway identity or synaptic placement. We first used multi-region single unit recordings during naturalistic behaviour and generalised linear models to estimate the functional connectivity from midbrain, hypothalamic, and cortical inputs onto dPAG neurons. We then combined synapse-resolution circuit tracing, two-photon dendritic stimulation with whole-cell somatic and dendritic recordings, and biophysical modelling to identify the mechanisms setting these weights. We found that dPAG neurons are electrotonically compact, generating broadly uniform somatic responses to inputs across the dendritic tree. As a result, presynaptic firing dynamics — burstiness within neurons and population synchrony — are the dominant determinants of input efficacy. This temporal-statistics framework accounts for the measured differences in functional connectivity across input regions and predicts that input weights should change dynamically whenever presynaptic temporal structure shifts — which we confirm by showing rapid, context-dependent reweighting of cortical input during motivational conflict. We propose that the subcellular specialisations of dPAG neurons allow them to integrate signals from distributed sources into a single decision, with input weights that can be flexibly adjusted on behavioural timescales — a principle that may extend to other brain hubs that compute survival decisions.

## Introduction

Escape from imminent threat is a fundamental survival behavior that requires rapid yet flexible decision-making. While escape is expressed by a well-defined repertoire of actions such as running or jumping, and coordinated autonomic responses such as tachycardia and hyperventilation, the decision of whether to escape is not a fixed stimulus-response reflex^1^. Rather, it reflects a flexible decision-making process that evaluates trade-offs between predation risk, defensive action outcomes, and the opportunity costs of forgoing other fitness-enhancing behaviours^2,3^. This requires integrating multiple external and internal factors such sensory stimulus properties^4,5^, environment structure^6^, prior experience^1,7,8^ and other survival needs such as hunger and thirst^1^. The final decision depends on the relative weight assigned to each of these information streams, which will change depending on the behavioural context.

Previous work has established the dorsal periaqueductal gray (dPAG) as the command center for escape initiation. Classical electrical and chemical stimulation and lesion studies showed that stimulation of dPAG evokes the full repertoire of escape-related behavioural and autonomic responses^9,10^ as well as subjective reports of panic in humans^11–13^, while dPAG lesions attenuate these responses^14,15^. Recent studies recording and manipulating the activity of molecularly defined populations of dPAG neurons have refined our understanding of their computational roles in escape^16–19^, positioning vGluT2+ dPAG neurons as gates that command the initiation and vigor of escape actions via integration and thresholding of upstream sensory threat evidence^4^.

Consistent with this gating role, the diverse information streams that shape escape decisions converge onto vGluT2+ dPAG neurons through inputs from multiple midbrain, hypothalamic and forebrain regions known to modulate escape behavior^15^. Sensory midbrain regions—the superior colliculus (SC) and inferior colliculus (IC)—provide acute information about visual and auditory threats, with SC-dPAG and IC-dPAG projections being necessary for visually and sound-evoked escape responses^4,20^. In contrast, dPAG-projecting regions in the medial hypothalamic defensive system, such as the ventromedial hypothalamus (VMH) and premammillary nucleus (PMd), integrate state variables over slower timescales to modulate escape^21–23^. Beyond threat-related information, dPAG neurons also receive additional modulatory inputs from the forebrain, most notably from the anterior cingulate cortex (ACC), which arbitrates competing threat, safety, and contextual signals to impose top-down ‘policy’ control^24–26^.

Despite this map of input pathways and their established effect on escape behaviour, it is unknown what determines their relative weights onto vGluT2+ dPAG neurons, and how they are integrated to compute escape decisions. On one hand, the influence of each pathway could be set by stable features of circuit architecture, such as differences in the number and density of synaptic contacts, their placement along the dendritic tree, or input-specific differences in synaptic strength. Such structural biases would impose a fixed baseline gain on each input, allowing, for example, acute threat signals from the sensory midbrain to be innately weighted more heavily than contextual signals from cortex. Alternatively, all inputs could share comparable baseline weights at the postsynaptic neuron, with their relative impact instead dominated by the activity dynamics of each presynaptic region. This arrangement would yield a more flexible system, in which input hierarchies could be rewritten on behaviourally relevant timescales by changes in upstream firing patterns and neuromodulation.

Here we have investigated how excitatory dPAG neurons integrate input from five major regions — SC, IC, VMH, PMd, and ACC — to compute escape decisions. We simultaneously recorded single-unit activity from dPAG and these input areas in freely behaving mice and used functional connectivity analysis to estimate the moment-by-moment influence of each region on dPAG spiking. To understand the mechanistic basis of input integration we then characterized dendritic integration properties, synaptic organization, and temporal statistics of presynaptic activity for each projection. Our results reveal that dPAG neurons act as electrotonically compact integrators, whose sensitivity to presynaptic dynamics rather than synaptic placement determines the functional impact of their inputs on spiking output. These findings provide a framework for understanding how distributed brain circuits converge onto a central escape hub and are integrated to dynamically compute flexible survival decisions.

## Results

### Quantifying the functional weight of long-range inputs onto dPAG neurons

To measure the functional connectivity strength of excitatory inputs from SC, IC, VMH, PMd, and ACC onto dPAG, we performed large-scale single unit recordings of these input regions and dPAG in freely behaving mice escaping from threat. We then fit generalised linear models^27–29^ (GLMs) to these data, in which the fitted parameters capture the coupling strength between each input and dPAG spiking. Simultaneous coverage of the input areas and dPAG was achieved using multiple chronically implanted, high density silicon probes (Neuropixels 2.0^30^), with one 4-shank probe placed posteriorly, passing through PAG, SC, IC and PMd, and one anteriorly, passing through ACC and VMH (Figure 1A; Figure S1 A, B). Across 6 mice and 26 recording sessions, we recorded a total of 4,480 units (496 PAG units, 482 SC units, 554 IC units, 152 VMH units, 63 PMd units and 2,733 ACC units), with an average of 19.1 ± 3.34 PAG units and 153.2 ± 11.4 units simultaneously recorded across the input regions in each session (Figure 1F, left; Figure S1C).

**Figure 1.**
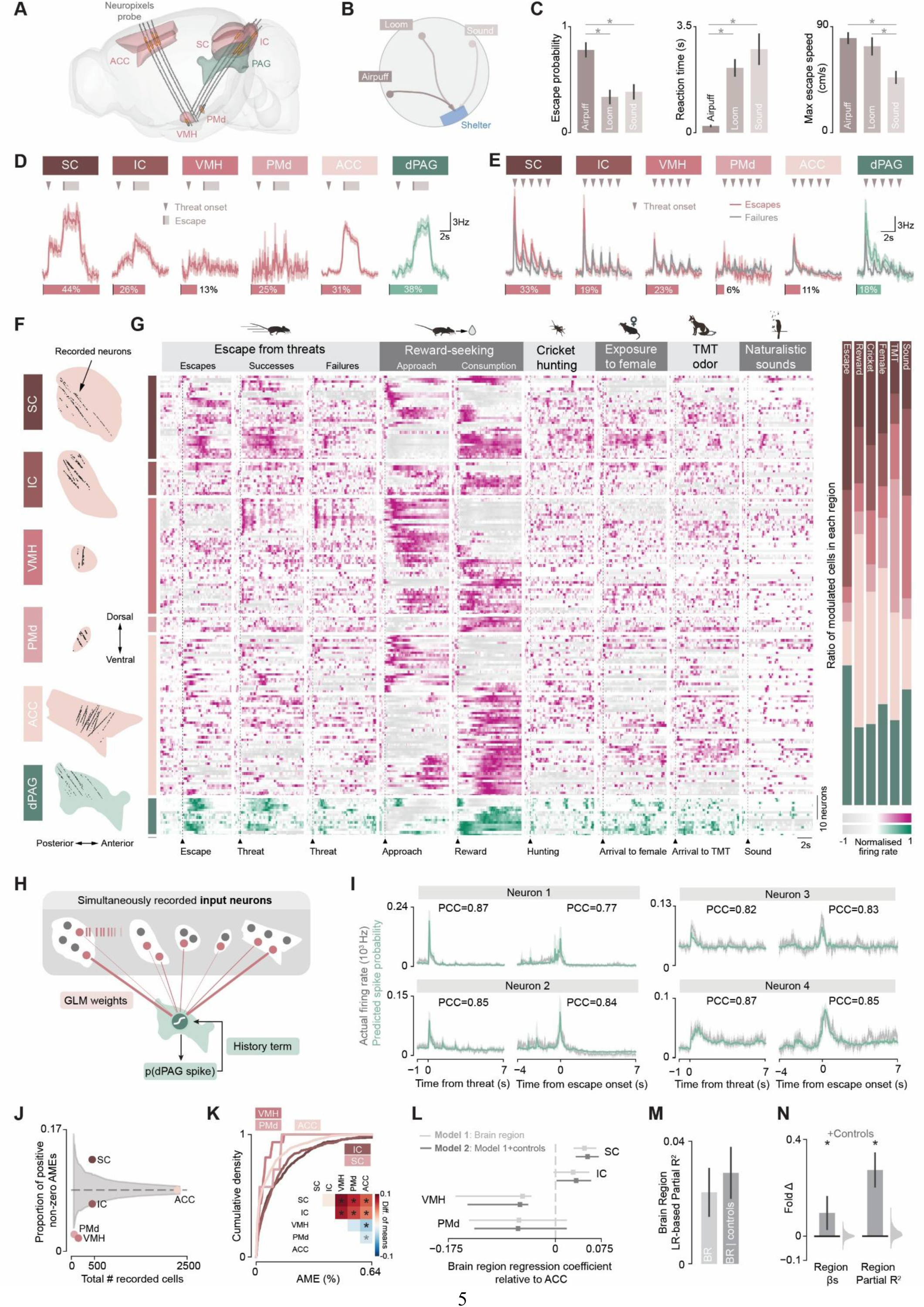
Regional differences in functional connectivity onto dPAG. **(A)** Schematic for simultaneous Neuropixels recordings from dPAG and its input regions: superior colliculus (SC), inferior colliculus (IC), ventromedial hypothalamus (VMH), dorsal premammillary nucleus (PMd), and anterior cingulate cortex (ACC). Grey lines show individual-shank trajectories from one example mouse; thicker portions indicate active recording sites in the session, with sites within target regions highlighted in yellow. **(B)** Example escape trajectories evoked by airpuff, visual loom, or auditory threat, from escape onset to shelter entry. **(C)** Session-averaged escape probability (left), trial-averaged reaction time (middle), and peak escape speed (right) for each stimulus type. Airpuffs evoked escapes with the highest probability (vs. loom, paired t-test p = 7.77 × 10⁻⁵; vs. sound, p = 1.92 × 10⁻⁴), shortest reaction times (vs. loom, permutation test p < 1 × 10⁻⁴; vs. sound, p < 1 × 10⁻⁴), and fastest peak speeds (vs. sound, permutation test p < 1 × 10⁻⁴; vs. loom, p = 0.817). Loom and sound evoked escapes at comparable probabilities (paired t-test p = 0.797) and reaction times (permutation test p = 0.972), but loom-evoked escapes reached higher peak speeds (permutation test p = 4 × 10⁻⁴). Error bars: parametric (left) or bootstrap-based (middle, right) SEM. **(D)** Average peri-escape PSTHs for escape-activated cells in each region (top), with the proportion of escape-activated cells indicated below. Activity on individual escape trials was time-warped to match escape and shelter entry latencies, then averaged across trials and aligned to stimulus onset (triangle), escape onset, and shelter entry (bar edges). **(E)** As in (D), but for cells activated by visual threat presentation, on escape trials (pink) and failure trials (grey). **(F)** Sagittal reconstruction of recorded single-unit locations across all sessions, by region. **(G)** Left: PSTHs from one example session for cells in each region, aligned to events across experimental blocks: escape onset and threat onset on success and failure trials (escape block); approach onset and reward consumption (reward block); contact with cricket, female mouse, or TMT-loaded filter paper (cricket, female, and TMT blocks); and onset of naturalistic sounds (sound block). Right: relative proportion of activity-modulated cells per block across regions, showing significant compositional variation across blocks (permutation-based test, p < 0.00001). **(H)** Schematic of Bernoulli GLM predicting dPAG spike probability from simultaneously recorded input spike trains. **(I)** Actual (grey) and predicted (green) peri-threat and –escape activity for four example dPAG cells. **(J)** Proportion of inputs with positive non-zero average marginal effect (AME, indicating excitatory functional connectivity). Grey shaded area indicates 1st-99th percentiles of null distribution under equal connectivity rate across all regions (grey dashed line), at each sampled population size. **(K)** Cumulative distributions of positive AME magnitudes for each region, in percentage points (pp) of dPAG spike probability per unit input. Inset: pairwise differences in mean AME (row − column); asterisks indicate significance (wild bootstrap test). Light grey asterisk: significant only with VMH and PMd pooled. **(L)** Regression coefficients and partial R² **(M)** of the brain-region factor in linear mixed models of AME magnitude, with random intercepts and either brain region alone (light grey) or brain region plus dPAG and input firing-rate controls (dark grey). Error bars: bootstrap SEM. **N)** Fold change in regional coefficients and partial R² with vs. without controls, showing that regional differences are not reduced when firing rates are accounted for. Light grey kernel-density estimates: null distribution from shuffled controls.

Mice were placed on an elevated open circular arena with a shelter at one edge, and presented with visual looming stimuli, airpuffs and auditory threats^31^ (Figure 1B, Figure S3A), which evoked escapes with varying probability, latency and vigour (Figure 1C). Neural activity during threat and escape was consistent with previous reports^4,20,22,23,25^, with dPAG responses dominated by a ramping increase in firing around escape onset. SC neurons showed a mixture of rapid, stimulus-locked responses that generalized across threat modalities and were stronger on escape trials, together with sustained activity during escape itself. IC neurons responded preferentially to auditory threats and were predominantly sensory, with little difference between escape and failure trials. VMH neurons were largely unresponsive during escape but showed transient, stimulus-locked responses at longer latencies than those observed in the midbrain. PMd neurons, by contrast, responded weakly to the threat stimulus but showed small, transient activity around escape onset and shelter entry. ACC neurons rarely showed threat-evoked responses, which when present were rapidly suppressed; activity instead increased just before escape onset. (Figure 1 D, E and Figure S2).

Since GLM inference requires sufficient variability in input and output activity to disentangle the independent contribution of each region to dPAG spiking, mice were also presented with an additional battery of ethological stimuli within the same recording session. To engage each input region, we used stimuli previously shown to drive its activity: live crickets (Figure S3B) to recruit SC during hunting^32,33^; a female conspecific (Figure S3C) to evoke IC responses to vocalisations^34^ and ACC responses to social signals^35^; predator odours (Figure S3E) to modulate VMH and PMd^15,22,36,37^; and naturalistic sounds (Figure S3D) to engage distributed auditory representations^38^ (Figure S3B). Mice were additionally trained on a food-seeking task expected to recruit ACC reward-seeking signals^39^ (Figure S3F-H).

This approach recruited distinct ensembles and activity profiles in neurons that might otherwise be coactive in a single behaviour (Figure 1G, left), with regions modulating their firing rates at differing ratios across blocks (permutation-based test of compositional differences, p < 0.00001; Figure 1G, right). This rich dataset of diverse, simultaneously recorded input and dPAG activity across naturalistic behaviors provided the foundation for estimating functional connectivity and predicting escape-related dPAG spiking. To quantify the relationship between input activity and dPAG spiking, we used Bernoulli GLMs, modelling the millisecond-wise spike probability in each dPAG cell as a function of the simultaneously recorded spike trains from all input regions^40–43^. This statistical framework has the advantage of capturing the point process nature of biological spikes—akin to soft-threshold leaky integrate-and-fire models^27,28^, while retaining the tractable likelihood-based inference and efficient parameter estimation of GLM formulation.

Inputs were convolved with an exponential kernel with a decay time constant matching the membrane time constant of dPAG cells (∼18 ms^28,44^), summed with varying weights to a membrane potential-like linear drive, and then mapped to spike probability via a logit link function (Figure 1H; see Methods). This yields coupling coefficients that serve as operational measures of influence conditional on the activity of all other simultaneously recorded neurons, in contrast to descriptive correlation methods, which are more strongly confounded by co-modulation and shared inputs^45^. We validated this approach by simulating feedforward circuits with known synaptic connectivity onto leaky integrate-and-fire neurons with experimentally derived parameters; the GLM successfully predicted postsynaptic spikes and recovered the underlying connectivity matrix, assigning near-zero weights to unconnected inputs and positive weights that scaled with synaptic conductance to excitatory synapses (Figure S4A).

We then fit GLMs to predict the activity of 496 recorded dPAG cells, and identified 114 cells whose peri-threat activity was predicted significantly above chance (permutation test, p<0.05, see Methods). The fitted activity profiles ranged from stimulus onset and longer lasting stimulus responses to escape-onset responses and showed good correspondence between recorded firing rate and predicted spike probability (average prediction accuracy across dPAG population – Pearson’s correlation coefficient: 0.570 ± 0.017) without requiring explicit behavioral information such as threat timing or locomotion speed (Figure 1I, Figure S2B). This suggests that our simultaneous recordings successfully captured cells in the input regions that are functionally connected to dPAG neurons. To compare coefficients across GLMs with different baseline firing probabilities and predictor sets, we converted raw GLM coefficients to average marginal effects (AMEs), expressing the effect of each input as the percentage point change in spike probability per presynaptic spike^46^ (Figure S4C). For subsequent analyses we focused on excitatory functional connections, identified from the distribution of AMEs (Figure S4C).

### Functional connectivity onto dPAG follows a regional hierarchy

The GLM-based analysis revealed that both the density and the strength of functional connections varied markedly across input regions (Figure 1J,K). Connection density, measured as the fraction of input cells significantly coupled to dPAG, was highest in the SC (12.8%), significantly above the overall average of 8.55% (permutation test, p < 0.001). IC and ACC showed intermediate densities (6.62% and 8.55%, respectively) not significantly different from average, while hypothalamic inputs were strikingly sparse: VMH and PMd connection densities were just 1.83% and 2.36%, well below null distributions accounting for their smaller sample sizes (permutation test, p < 0.001 and p = 0.007, respectively; Figure 1J). The coupling strength also differed significantly across regions (bootstrap ANOVA, F = 18.9, p < 0.00001; Figure 1K). Within each anatomical class, member regions were comparable: SC and IC did not differ from one another, nor did VMH and PMd. Across classes, however, midbrain inputs exerted the largest influence on dPAG spike probability (SC: 0.133 ± 0.008; IC: 0.124 ± 0.010 percentage points), approximately 1.8-fold greater than ACC (0.073 ± 0.004 pp; bootstrap test, p < 0.00021 for both) and 3-fold greater than hypothalamic inputs (VMH: 0.040 ± 0.009 pp; p < 0.00035; PMd: 0.057 ± 0.022 pp; p < 0.027). Hypothalamic inputs had the weakest influence of the three classes overall (p < 0.009 for all comparisons between pooled VMH and PMd data and other regions).

While dPAG contains both excitatory and inhibitory neurons^47,48^, glutamatergic cells are almost three times as numerous as GABAergic cells^48^ and have larger cell bodies^48^, and thus most dPAG units in our recordings are likely excitatory. Nevertheless, to confirm that the same input hierarchy holds for genetically defined excitatory dPAG neurons, we expressed Cre-dependent ChR2 in dPAG of vGluT2–Cre mice (Figure S5A) and optotagged the labelled cells during recording with blue-light stimulation (Figure S5B; n = 28 units). Both connection density and strength from each input region onto vGluT2+ neurons matched those measured across the full dPAG population (Figure S5C), indicating that the regional pattern is intrinsic to glutamatergic dPAG neurons. To further confirm that the same input hierarchy holds for anatomically verified dPAG-projecting neurons, rather than reflecting incidental coupling to neighbouring or passing cells, we delivered retrograde AAV to the dPAG to drive ChrimsonR expression in dPAG-projecting input neurons (Figure S5A) and identified them online via antidromic spikes evoked by red-light stimulation at dPAG (Figure S5D; n = 121 units). As expected, connection densities among these anatomically verified projection neurons were elevated relative to the general population (1.7-, 7.8-, and 1.4-fold for ACC, SC, and VMH respectively). Critically, however, both the rank order across regions and the relative connection strengths were preserved (Figure S5E), confirming that the regional differences are a property of the dPAG-projecting populations themselves.

Finally, we considered whether regional differences could arise from systematic biases in firing rates, since low firing rates in inputs and dPAG cells yield underestimated coefficients due to sparse informative bins (Figure S4D, E). We therefore fit linear mixed-effect models (LMMs) that included input and dPAG time-averaged firing rates as control variables. Brain region held significant explanatory power for AME magnitude s (likelihood-ratio test, p = 1.08 × 10⁻⁷), and inclusion of these variables did not reduce its explanatory power nor coefficients (Figure1L-N). Thus, the regional differences in functional connectivity cannot be explained by differences in firing rates and reflect genuine differences in the strength of influence that each input region exerts on dPAG spiking.

Together, these results reveal that in the behavioural context analysed there is a hierarchy of functional connectivity to dPAG: midbrain sensory regions (SC and IC) show both the highest connectivity rates and the strongest individual connections, cortical input (ACC) shows intermediate connectivity, and hypothalamic regions (VMH and PMd) have the weakest and sparsest functional coupling to dPAG.

### Synaptic organization of inputs onto dPAG does not predict their functional weight

We next sought to understand the mechanisms underlying these regional functional connectivity differences. First, we considered that biases in synapse location could be a candidate mechanism since it can critically shape the neuronal input-output function through electrotonic filtering and differential engagement of active conductances^49–51^. We therefore generated subcellular maps of synapse locations for glutamatergic inputs from SC, IC, VMH, PMd, and ACC onto glutamatergic dPAG neurons.

To label excitatory presynaptic terminals, each input region was separately injected with an AAV encoding Cre-dependent synaptophysin-mCherry in vGluT2-Cre mice (Figure 2A, S6A). Tracing the long dendrites of dPAG neurons without interference from neighboring cells required sparse labelling of postsynaptic dPAG neurons. We achieved this with an anterograde transsynaptic strategy (Figure 2A; see Methods): AAV2/1-flexed-Flp was co-injected at the input region to exploit the inefficiency of anterograde transsynaptic jumping^52^ and drive sparse Flp expression in vGluT2+ dPAG neurons; and a second AAV was injected at dPAG, carrying a Flp-dependent, TRE-tTA-autoregulated GFP construct. To image synapse location we cleared thick coronal sections encompassing dPAG using CUBIC^53^, which enabled confocal imaging depths up to 2,288 μm. We then reconstructed dendritic trees in 3D^54^ and identified putative synapses by colocalization of mCherry and GFP signal, with manual curation in 3D to confirm close apposition of presynaptic terminal against dendrite (Figure 2B). This yielded subcellular maps of 1,715 synapses onto 97 dPAG neurons across the five input regions (Figure 2C).

**Figure 2.**
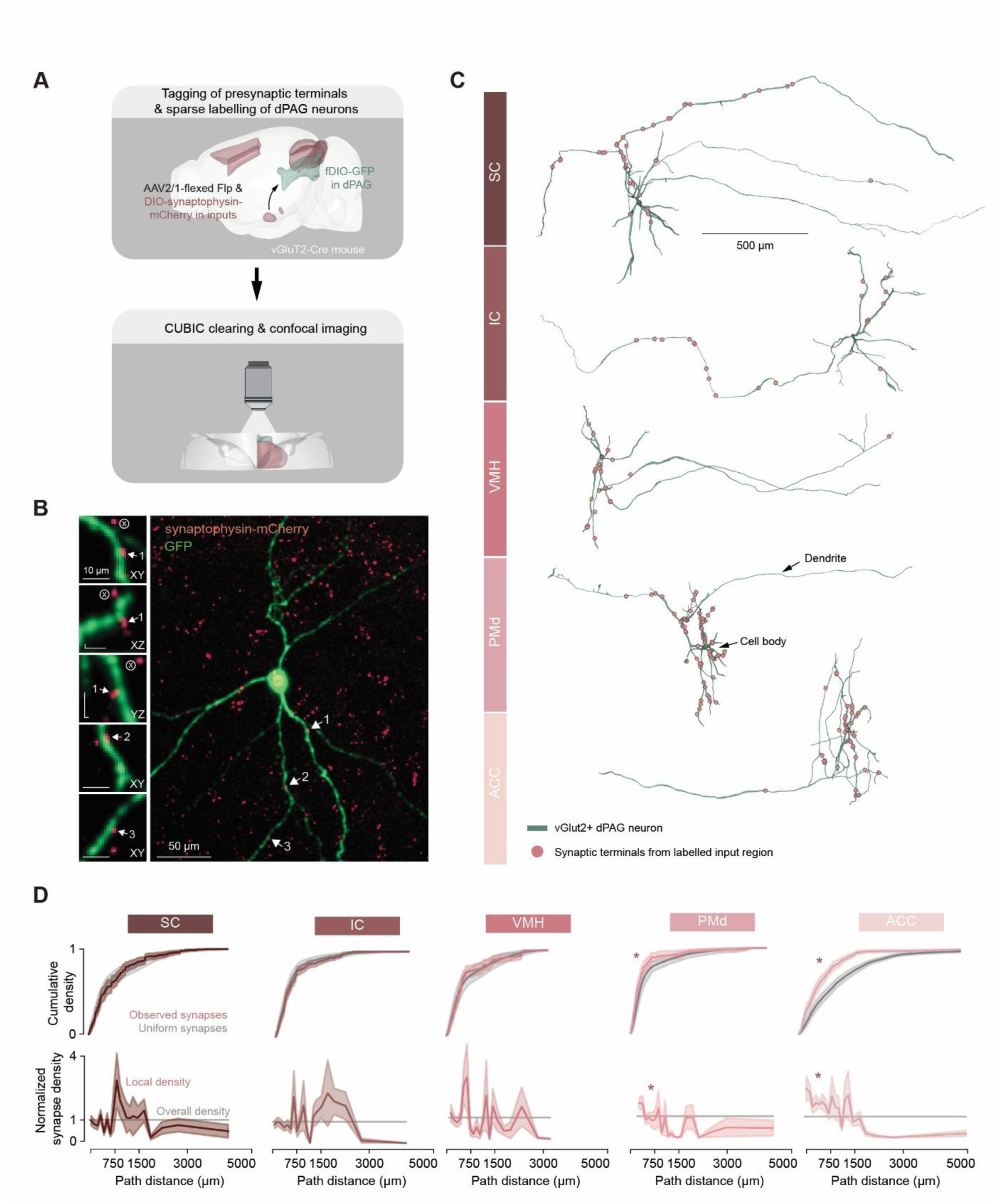
Synaptic organisation of long-range inputs onto dPAG dendrites. **(A)** Viral and imaging strategies for mapping excitatory presynaptic terminals from each input region onto vGluT2+ dPAG neurons. **(B)** Example showing GFP-labelled dPAG dendrites and synaptophysin-mCherry-labelled presynaptic terminals. Arrows mark putative synaptic contacts, shown at higher magnification on the left. Synapse 1 is shown in three orthogonal views to confirm close apposition of the presynaptic terminal to the dendrite. The circled “×” marks an mCherry puncta close to the dendrite in some views only and therefore not classified as a synapse. **(C)** Example 3D reconstructions of dPAG neurons (green) and their synapses (red spheres) from each input region. **(D)** Top: cumulative distributions of observed synapse path distances to soma, averaged across cells (pink, with SEM shading), compared with the null distribution of independent uniform placement along the traced dendrite (grey, with 1st–99th-percentile shading). Asterisks: significant deviation from uniform (bootstrap-based KS test). Bottom: local synapse density as a function of path distance, normalised to the overall synapse density. Asterisks: significantly non-constant density (bootstrap-based KPSS test).

We first quantified synapse path distance to soma—the cumulative length along the dendritic tree between each synapse and the soma (Figure 2D)— which is a key determinant of synaptic integration^55^. To account for differences in traced dendrite lengths across neurons, we compared synapse distributions against per-neuron null distributions generated by uniform resampling along the dendritic tree. SC, IC and VMH synapses were uniformly distributed along the dendritic tree, with path distance distributions not different from chance (bootstrap-based Kolmogorov-Smirnov test, p = 1 for each; Figure 2D). In contrast, ACC and PMd synapses were significantly enriched at proximal locations closer to the soma (p < 0.0001 and p = 0.005 respectively). This translated to significant variation of synapse density with path distance for ACC and PMd inputs (bootstrap-based KPSS test for stationarity, p = 0.018 for both), but uniform density for SC, IC and VMH inputs (p = 1 for each; Figure D).

We next examined spatial relationships between synapses using nearest-neighbor distance as a measure of clustering (Figure S6B). SC and ACC synapses showed significantly smaller nearest-neighbor distances than expected for uniform distribution across branches (bootstrap-based KS test, p < 0.0001 and p = 0.004 respectively), indicating a tendency for clustering on the same or nearby dendritic segments. In contrast, IC, VMH and PMd synapses showed nearest-neighbor distances consistent with dispersed placement across different branches (p = 1, 1 and 0.513 respectively).

Together, these data reveal a largely uniform distribution of synaptic inputs onto dPAG dendrites, with only a few region-specific deviations: proximal enrichment of ACC and PMd synapses, and clustering of SC and ACC synapses. These deviations do not, however, map onto the observed differences in functional connectivity. Proximal enrichment might be expected to enhance efficacy through reduced dendritic filtering^49–51,55^, yet the proximally enriched PMd inputs are the weakest of all five regions. Likewise, while clustering can in principle either enhance or attenuate efficacy depending on the integration regime^56,57^, SC inputs show clustering yet are no different than the dispersed IC inputs. Taken together, these dissociations indicate that synapse placement alone cannot account for the regional differences in functional connectivity, pointing instead to dendritic biophysical properties or the temporal statistics of presynaptic activity as the more likely determinants of input efficacy.

### dPAG dendrites support broadly uniform synaptic integration

While dendritic biophysical properties have been extensively studied in cortical neurons^58^, signal propagation in dPAG dendrites has not been previously measured. Given that these dendrites are long and thin^59–61^, we initially expected significant electrotonic attenuation of distal inputs^62,63^, comparable to that seen in other fine dendrites such as the basal dendrites of layer 5 pyramidal neurons^64^ and dendrites of dentate gyrus granule cells^65,66^.

To test this, we used *in vitro* two-photon holographic stimulation to activate ChR2-expressing vGluT2+ dPAG neurons at defined dendritic locations while recording somatic voltage responses with whole-cell patch clamp (Figure 3A; see Methods). Across 38 dPAG cells, we sampled 618 dendritic sites at distances up to 554 μm from the soma, all of which evoked detectable somatic responses. We quantified signal propagation using the rise time of the somatic response, which is largely invariant to differences in ChR2 expression and dendritic diameter — factors that produce variability in photocurrent amplitude and would otherwise confound amplitude-based measurements (Figure S7; see Methods). We found that rise times increased modestly with dendritic distance, consistent with capacitive filtering of distal inputs (Figure 3B-D). However, the degree of temporal filtering was markedly smaller than expected: dPAG dendrites filtered at 9.20 ± 0.68 ms/mm, less than half the rate in the similarly fine dendrites of dentate gyrus granule cells (DG GC; 20.7 ± 0.16 ms/mm in simulations; Wald test, p < 0.0001; Figure 3D, K), suggesting that dPAG are surprisingly electrotonically compact.

**Figure 3.**
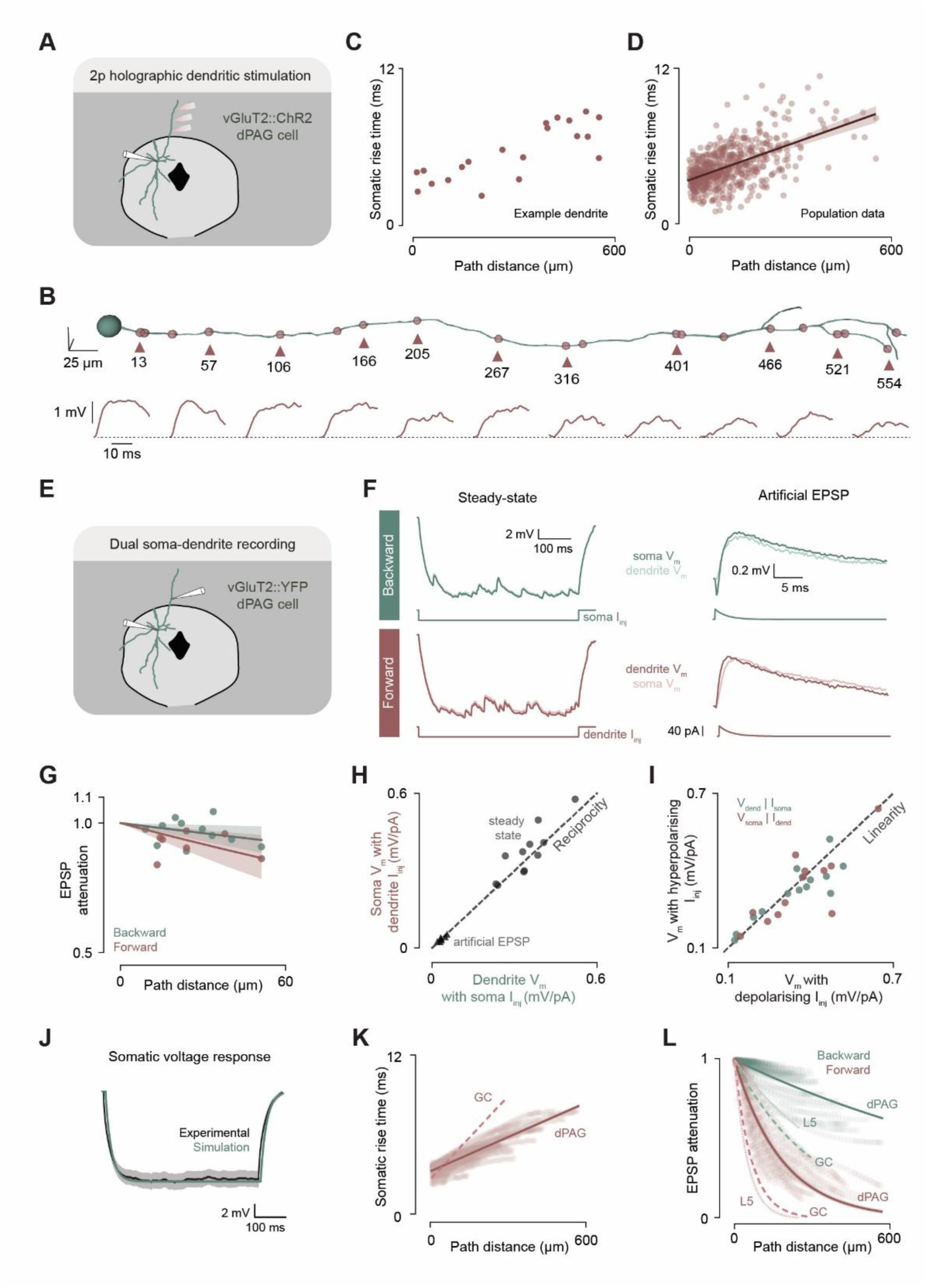
vGluT2+ dPAG dendrites are electrotonically compact. **(A)** Schematic of two-photon holographic stimulation of ChR2-expressing dendrites of vGluT2+ dPAG neurons *in vitro*, with whole-cell recording at the soma. **(B)** Top: example ChR2-expressing dendrite (green) and stimulation points (dark pink), with path distances to the soma indicated (in μm). Bottom: somatic responses to stimulation at the locations marked with triangles. **(C-D)** Rise time of somatic responses for the example dendrite in B (C) and pooled across all dendrites (D). **(E)** Schematic of simultaneous somatic and dendritic whole-cell recording in vGluT2+ dPAG neurons *in vitro*.**(F)** Example voltage responses at the soma and dendrite to hyperpolarising step (left) and depolarising EPSC-like (right) current injections delivered at the soma (top, green) or dendrite (bottom, pink). **(G)** Backward (green; soma → dendrite) and forward (pink; dendrite → soma) attenuation of artificial EPSP responses, quantified as the ratio of voltage at the recording-only site to voltage at the stimulation site. Effective length constants: 754 ± 292 μm backward (n = 11) and 352 ± 86 μm forward (n = 7). **(H)** Reciprocity of soma–dendrite input transfer, quantified as the ratio of soma response to dendritic injection over dendrite response to somatic injection, for steady-state (circles; 0.948 ± 0.038; 1-sample t-test vs. 1, n = 11, p = 0.20) and artificial EPSP (triangles; 1.019 ± 0.050; n = 7, p = 0.72) responses. **(I)** Linearity of input transfer for steady-state responses, quantified as the ratio of recording-site response to hyperpolarising vs. depolarising step current, in the backward (green; 0.925 ± 0.147; n = 12, p = 0.62) and forward (pink; 1.13 ± 0.089; n = 14, p = 0.17) directions. P-values from 1-sample t-tests against 1. **(J)** Experimentally measured (black) and simulated (green; passive multi-compartmental model) somatic voltage responses to hyperpolarising step current injection. **(K)** Somatic rise times in response to simulated ChR2 photostimulation along dPAG dendrites (dark pink), showing a shallower relationship with path distance than dentate gyrus granule cell (GC) models (dashed line, light pink). **(L)** Simulated backward (green) and forward (pink) attenuation of artificial EPSP responses in dPAG models (solid lines), shown alongside dentate gyrus granule cell dendrites (GC, dashed lines) and layer 5 pyramidal cell basal dendrites (L5, dotted lines). GC values from Schmidt-Hieber et al., 2007 and Krueppel et al., 2011; L5 values from Nevian et al., 2007.

To measure signal attenuation directly, we next performed dual somato-dendritic whole-cell recordings in vGluT2+ dPAG neurons *in vitro* (Figure 3E). Although only proximal dendrites could be reliably targeted, injection of steady-state current steps and EPSC-like transients (Figure 3F) revealed highly efficient electrotonic coupling between soma and dendrite (Figure 3G and S8C). Effective length constants for forward EPSP propagation (352 ± 86 μm) and backward steady-state propagation (1,403 ± 446 μm) were significantly longer than those measured in layer 5 pyramidal cell basal dendrites or DG granule cell dendrites (Figure 3L and S8D; Wald test, p < 0.01 for all comparisons). Transfer impedance showed reciprocity and linearity, consistent with passive propagation in the subthreshold range at these distances (Figure 3H and I). Input resistance was high both at the soma (352 ± 32 MΩ, Figure S8A) and dendrite (399 ± 32 MΩ), substantially exceeding typical values for cortical and hippocampal pyramidal neurons *in vitro*^67^. Together, these direct measurements further support the conclusion that dPAG dendrites are electrotonically compact.

To test whether this electrotonic compactness could be explained by passive cable properties alone, we constructed multi-compartmental biophysical models using experimentally reconstructed dPAG morphologies, with specific membrane resistance and axial resistance as free parameters (see Methods). Passive cable models successfully reproduced all experimental observations—input resistance, membrane time constant (Figure 3J, S8A-B), forward and backward attenuation of steady-state (Figure 3G & L) and transient signals (Figure S8C-D), and distance-dependent rise times (Figure 3D & K)—without requiring voltage-gated conductances or non-uniform membrane properties. The fitted axial resistance (105–125 Ω·cm) was lower than values reported for DG granule cells or CA1 pyramidal neurons^65,68^, and dPAG dendrites were sparsely branched relative to DG granule cells (15.6 ± 1.4 vs 32.3 ± 2.3 branches per cell), with both features contributing to the observed compactness.

We next used these experimentally constrained models to investigate synaptic integration in dPAG neurons, comparing the results to equivalent models of DG granule cells. We first simulated activation of individual synapses across the dendritic tree (Figure 4A) and measured somatic response amplitudes (Figure 4B). In dPAG neurons, somatic responses remained relatively constant across synapse locations (length constant 809 ± 11 μm, Figure 4C, left), whereas in DG granule cells they decayed steeply with distance (length constant 227 ± 1 μm; Figure 4C, right). This near-invariance substantially reduced the boosting of somatic responses expected from proximal synapse enrichment: given the path distance distributions measured anatomically, PMd and ACC synapses should produce somatic responses only 8% and 51% larger than uniformly distributed synapses in dPAG neurons, compared to 33% and 86% larger in DG granule cells (Figure S10A).

**Figure 4.**
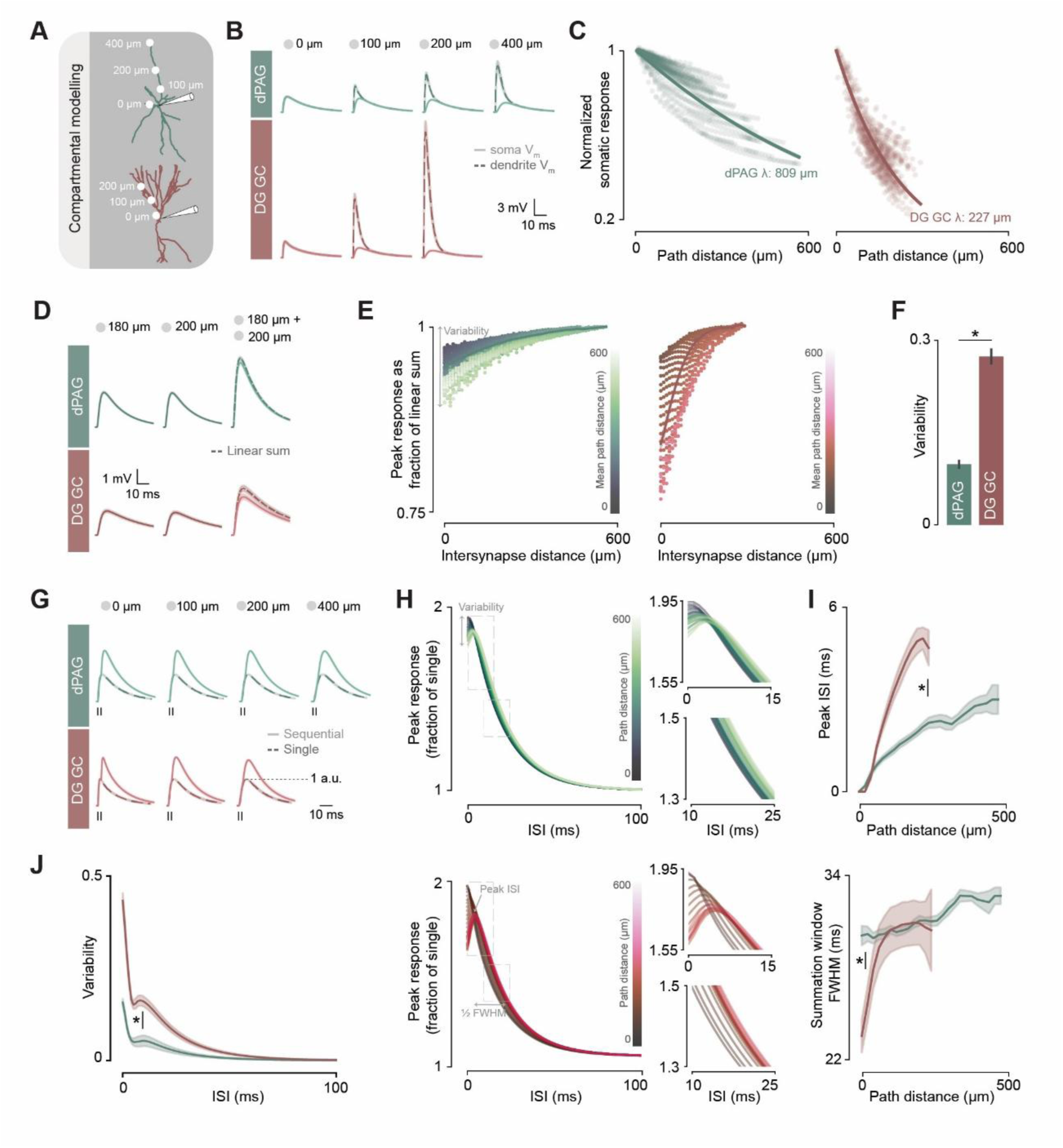
Weakly location-dependent synaptic integration in vGluT2+ dPAG neurons. **(A)** Compartmental models of vGluT2+ dPAG (green) and dentate gyrus granule cell (DG GC; pink) neurons, with synapse locations used in simulations marked (white circles). **(B)** Example voltage responses at the soma (light solid lines) and at the local dendritic site (dark dashed lines) to synaptic stimulation at different path distances, for dPAG (top, green) and DG GC (bottom, pink). Somatic responses diminish with path distance in DG GC but remain relatively stable in dPAG. **(C)** Peak somatic response, normalised to the response evoked by the most proximal synapse, as a function of synapse path distance for dPAG (left) and DG GC (right). **(D)** Example voltage responses to stimulation of each of two synapses individually (left, middle) and simultaneously (right; light solid lines), overlaid with the linear sum of the individual responses (dark dashed lines). Spatial summation is the ratio of the peak simultaneous response to the peak linear sum. **(E)** Spatial summation as a function of inter-synapse distance and average synapse path distance from the soma (colour intensity), for dPAG (left) and DG GC (right). Each point is the mean within a 10 μm × 10 μm bin of inter-synapse distance and average path distance. **(F)** Average within-cell variability (min–max range) in spatial summation, for dPAG and DG GC. **(G)** Example voltage responses to sequential synaptic stimulation at 2 ms interstimulus interval at varying path distances (light solid lines), overlaid with responses to single stimulation of the same synapse (dark dashed lines). Responses are normalised to the peak of the single-stimulation response. Temporal summation is the ratio of the peak sequential response to the peak single-stimulation response. **(H)** Temporal summation as a function of interstimulus interval and synapse path distance (colour intensity, curves binned by path distance in 10 μm bins), for dPAG (top, green) and DG GC (bottom, pink). **(I)** Variation of the temporal summation curve with synapse path distance, characterised by the interstimulus interval at which summation peaks (top) and the width of the summation window (bottom). **(J)** Average within-cell variability (min–max range) in temporal summation, for dPAG (green) and DG GC (pink), at each interstimulus interval.

We next examined spatial summation by co-activating pairs of synapses at varying dendritic locations. In both dPAG and DG granule cell models, summation was sublinear when synapses were close together, due to driving force saturation (Figure 4D) – more strongly for synapses on the same branch (Figure S10B). However, the variability in spatial summation across all possible synapse pairs was substantially lower in dPAG neurons (range: 0.098 ± 0.006) than in DG GC (range: 0.274 ± 0.011; t-test, p < 0.0001; Figure 4E, F), indicating that spatial summation in dPAG is relatively insensitive to the precise configuration of co-active synapses.

Finally, we examined temporal summation by activating individual synapses twice at varying interstimulus intervals. Temporal summation peaked at short intervals, when responses to successive inputs overlapped, and was reduced at near-simultaneous intervals at distal locations due to local driving force saturation (Figure 4G). In dPAG neurons, the shape of the temporal summation curve (Figure 4H, top) —including its peak interval and width (Figure 4I)—varied minimally with synapse location. In DG granule cells, by contrast, the curve showed pronounced location-dependence (Figure 4H, bottom), reflecting greater capacitive filtering of distal inputs. As a result, variability in temporal summation across all possible synapse locations was much lower in dPAG neurons than in DG granule cells at every interstimulus interval tested (t-test, p <= 0.0176 for all interstimulus intervals; Figure 4J).

Together, these experimental and modelling data suggest that dPAG neurons are electrotonically compact, with dendritic properties that render somatic responses only weakly sensitive to synapse location. This compactness minimises the influence of synapse placement on input efficacy, providing a mechanistic account for the dissociation between synapse organisation and functional connectivity reported above, and indicating that dPAG neurons integrate their inputs in a manner approximating a leaky integrate-and-fire point neuron rather than exploiting location-dependent dendritic computations. The regional differences in functional connectivity must therefore arise not from where inputs are placed on the dendritic tree, but from how presynaptic activity is structured in time.

### Presynaptic temporal statistics explain regional differences in functional connectivity

Having ruled out synapse placement as the source of regional differences in functional connectivity, we considered that the key determinant of input influence might instead be presynaptic activity dynamics^69,70^. Tightly timed presynaptic spikes summate more effectively than temporally dispersed inputs^50,51,71^, and therefore the impact of a presynaptic spike on dPAG spiking probability should depend on the temporal context of ongoing synaptic activity.

We first examined the temporal structure of individual input neurons by analyzing their interspike interval (ISI) distributions (Figure S11A). Each neuron’s ISI distribution was modelled as a gamma renewal process with shape parameter α: values near 1 indicate Poisson-like firing, α > 1 indicates regular pacemaker-like firing, and α < 1 indicates irregular firing with clustered short ISIs^72,73^ (Figure 5A). We found that the shape parameter differed significantly across regions (bootstrap-based ANOVA, p < 0.00001; Figure 5B and S11B, top): SC (0.652 ± 0.026), IC (0.652 ± 0.023) and ACC (0.525 ± 0.005) were all significantly lower than VMH (0.763 ± 0.032) and PMd (0.814 ± 0.035; bootstrap-based test of difference of means, p < 0.006 for all comparisons). SC, IC and ACC neurons therefore fire more irregularly than VMH and PMd neurons, with an excess of short ISIs. To quantify the expected postsynaptic consequence of this irregularity, we convolved each neuron’s ISI distribution with the temporal summation curve derived from our dPAG biophysical models and compared the predicted response size to that of matched Poisson spike trains (Figure 5C). Because spike trains enriched in short ISIs produce greater temporal summation, SC, IC and ACC inputs showed larger response boosting over their Poisson equivalents (4.2 ± 0.3%, 5.8 ± 0.3% and 6.5 ± 0.1% respectively) than VMH and PMd inputs (1.6 ± 0.4% and 2.1 ± 0.6%; bootstrap-based test of difference of means, p < 0.002 for all comparisons; Figure 5D and S11B, bottom).

**Figure 5.**
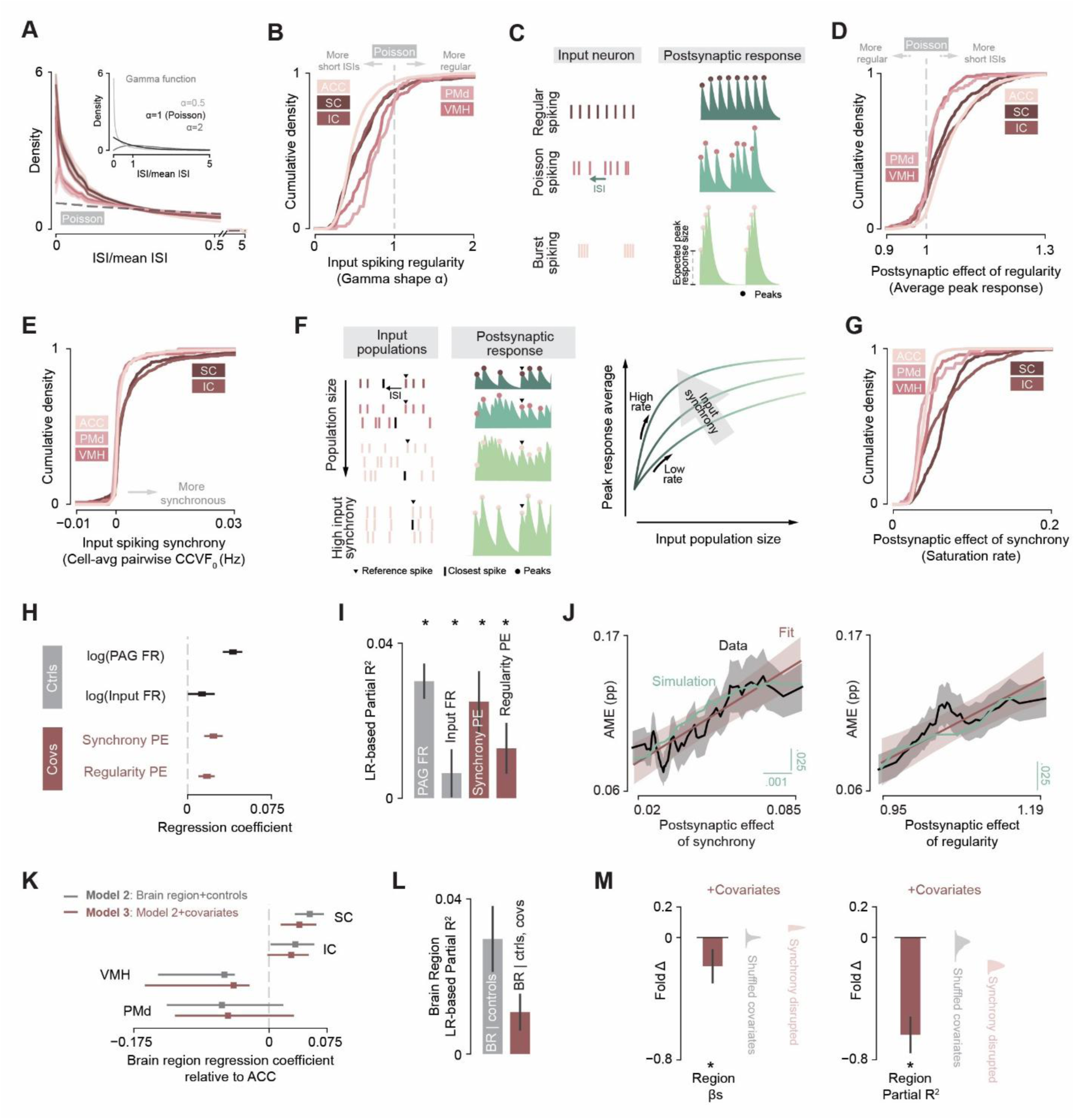
– Presynaptic temporal dynamics shape functional connectivity onto dPAG. **(A)** Region-averaged distributions of mean-normalised inter-spike intervals (ISI). The shape of the distribution reflects spiking regularity independently of firing rate and can be modelled by a gamma distribution with shape parameter α (inset). **(B)** Cumulative distributions of α across brain regions. **(C)** Schematic of postsynaptic temporal summation. The same number of presynaptic spikes at different regularities (left) produces different postsynaptic membrane potentials (right): peak responses (circles) are larger for presynaptic trains with higher irregularity and shorter ISIs (bottom). The mean peak response can be approximated by convolving the postsynaptic summation curve with the presynaptic ISI distribution. **(D)** Cumulative distributions of mean peak postsynaptic responses normalised to those evoked by Poisson spike trains at the same firing rate. This serves as an index of the postsynaptic effect of presynaptic regularity. **(E)** Cumulative distributions of cell-averaged pairwise cross-covariance at 0 ms lag (CCVF₀), reflecting the average rate of coincident spikes between a cell and others in the same region. **(F)** Schematic of postsynaptic temporal summation across multiple presynaptic cells. Left: peak postsynaptic responses (circles) evoked by a fixed set of spikes from one presynaptic cell (first cell in each population) increase as more presynaptic cells converge onto the same postsynaptic cell, because spikes from the focal cell (arrow) are more likely to fall within a short interval after a spike from another cell in the population (black). This effect is amplified when the focal cell’s spikes are synchronous with the rest of the population (bottom). Right: mean peak response increases with population size, at a higher rate for higher synchrony. **(G)** Cumulative distributions of the population saturation rate of mean peak postsynaptic response, an index of the postsynaptic effect of synchrony. **(H-I)** Regression coefficients (H) and likelihood-ratio-based partial R² (I) of the fixed effects in a linear mixed model of AME magnitude with random intercepts. Fixed effects include firing-rate controls (dPAG and input rates; grey) and the temporal-statistics covariates (Poisson-normalised mean responses, *regularity postsynaptic effect*; population saturation rates, *synchrony postsynaptic effect*; red). Error bars: bootstrap SEM. **(J)** Partial residuals (black, binned means) for synchrony (left) and regularity (right) postsynaptic effects after removing the contribution of other fixed and random effects, with liner mixed model predictions (red) and AMEs from simulated spike trains (green) overlaid. Shaded areas: 95% confidence intervals. **(K-L)** Regression coefficients of brain-region contrasts (K) and likelihood-ratio-based partial R² of the brain-region factor (L) in linear mixed models of AME magnitude, with random intercepts and either brain region with firing-rate controls (grey) or brain region with controls and temporal-statistics covariates (red) as fixed effects. Error bars: bootstrap SEM. **(M)** Fold change in brain-region coefficients (left) and partial R² (right) upon adding the temporal-statistics covariates. Light grey kernel-density estimates: null distribution from shuffled covariates. Light pink: null distribution from circular-shifted spike trains, which preserve firing rate but disrupt synchrony. Error bars: bootstrap SEM.

We next examined temporal relationships *across* neurons, since postsynaptic summation is agnostic of presynaptic source. We quantified pairwise synchrony as the cross-covariance at zero lag, which reflects spike coincidences above chance^74^. Synchrony differed significantly across regions (bootstrap-based ANOVA, p < 0.00001): SC (3.96 × 10⁻³ Hz) and IC (5.81 × 10⁻³ Hz) were significantly more synchronous than VMH (1.41 × 10⁻³ Hz), PMd (0.93 × 10⁻³ Hz) and ACC (1.52 × 10⁻³ Hz; bootstrap-based test of difference of means, p < 0.002 for all comparisons; Figure 5E and S11F, top). To estimate the impact of this synchrony on postsynaptic summation, we computed how the postsynaptic responses increased as spikes from additional neurons in the same region were progressively included. As population size grows, spikes become more likely to fall within the temporal summation window, until the response saturates; in synchronous populations, this saturation occurs more rapidly (Figure 5F and S11C-E). The saturation rate also differed significantly across regions, with SC (0.065 ± 0.001) and IC (0.062 ± 0.002) significantly higher than VMH (0.045 ± 0.002), PMd (0.042 ± 0.003) and ACC (0.038 ± 0.0002; bootstrap-based test of difference of means, p < 0.0002 for all comparisons; Figure 5G and S11F, bottom).

Together, these analyses reveal regional differences in both within-neuron regularity and population synchrony that translate into corresponding differences in postsynaptic temporal summation in dPAG neurons. These differences would be expected to produce stronger functional connectivity for midbrain inputs, and weaker connectivity for hypothalamic inputs — matching the pattern we observed *in vivo*. To test this relationship directly, we added per-neuron estimates of the postsynaptic effect of regularity and synchrony as covariates to the LMM of AME magnitude. Both covariates received significantly positive coefficients (permutation test, p < 0.001, Figure 5H), held significant explanatory value (likelihood-ratio test, p = 1.75 × 10⁻¹⁰ and 4.77 × 10⁻⁶ respectively; Figure 5I), and showed a linear relationship with AME after accounting for other covariates (Figure 5J). To establish causality, we parametrically increased presynaptic firing irregularity and synchrony in simulated feedforward networks: both manipulations increased GLM-estimated functional connectivity strength (Figure 5J and S12A-D), confirming that input temporal statistics are causal determinants of functional connectivity in this framework.

Finally, we asked whether these temporal statistics could explain the regional functional connectivity differences observed *in vivo*. Including the temporal statistic covariates alongside brain region identity in the LMM substantially reduced the contribution of brain region identity itself: region contrast coefficients of decreased by 18.8 ± 11.5% (permutation test, p < 0.001; Figure 5K), and overall explanatory value of brain region decreased by 63.6 ± 12.7% (permutation test, p < 0.001; Figure 5L). These reductions were significantly larger than those obtained with shuffled covariates (permutation test, p < 0.001; Figure 5M), indicating that temporal statistics captured variance genuinely attributable to brain region rather than a modelling artifact. As a further control, we recomputed the synchrony covariate from circularly shifted spike trains, which preserved firing rates and chance coincidences but disrupted true inter-neuronal synchrony. This control covariate showed substantially reduced explanatory contribution (Figure 5M), confirming that the effect of synchrony was driven by structured temporal coordination rather than rate alone.

Together, these results show that systematic regional differences in presynaptic temporal statistics largely account for the regional differences in functional connectivity to dPAG. Strong SC and IC connectivity reflects both higher within-neuron spike irregularity and higher population synchrony – features that favor temporal summation in electrotonically compact dPAG neurons. The weaker functional connectivity of VMH and PMd, in contrast, reflects more Poisson-like firing and lower synchrony, which together produce temporally dispersed inputs that summate less effectively.

### ACC influence on dPAG is dynamically reweighted during motivational conflict

Our findings suggest that the functional connectivity between input regions and dPAG neurons could vary dynamically depending on the temporal structure of the inputs. To test this, we asked whether ACC-dPAG functional connectivity changes across behavioural contexts in a manner explainable by changes in ACC temporal firing statistics. We focused on ACC because of its established role in evaluating competing drives^24^ and designed a behavioural manipulation expected to engage it selectively. We extended the food-seeking assay described above (Figure S3F-H) by introducing threatening stimuli on a subset of trials, creating a state of motivational conflict in which mice were required to weigh reward against safety. This allowed us to compare ACC firing patterns and ACC-dPAG functional connectivity across two conditions: *threat only blocks*, in which decisions could be guided by threat information alone, and *conflict blocks*, in which decisions required balancing threat against reward and were therefore expected to engage ACC more strongly.

When reward was available, mice expressed behavioural patterns consistent with active motivational conflict, evident in modulations of both reward-seeking (Figure 6A-E) and escape (Figure 6F-G) behaviours. Threats presented *en route* to the lick port produced a range of trajectory changes, from outright abortion of the approach to subtle initial reorientations toward the shelter that were subsequently overridden by a return to the lick port (Figure 6A, F). These responses jointly reduced the proportion of rewarded approaches (Figure 6B; 41.9 ± 5.63 % vs 79.3 ± 2.43 %; paired t-test, p = 8.93 x 10^−6^) and increased the time taken to reach the port (Figure 6C; 6.35 ± 1.14 s vs 2.53 ± 0.205 s; permutation test, p < 0.0002). Threats presented on arrival at the lick port caused mice to leave earlier (Figure 6D; 5.52 ± 0.874 s vs 7.80 ± 0.761 s; permutation test, p < 0.0004), and often to forgo the reward altogether (Figure 6B; 60.3 ± 6.66 % vs 79.3 ± 2.43 %; paired t-test, p = 8.94 x 10^−3^); threats presented just after mice left the port redirecting them to the shelter and prolonged the time to the next approach (Figure 6E; 38.1 ± 8.97 s vs 17.1 ± 1.31 s; permutation test, p = 0.0006).

**Figure 6.**
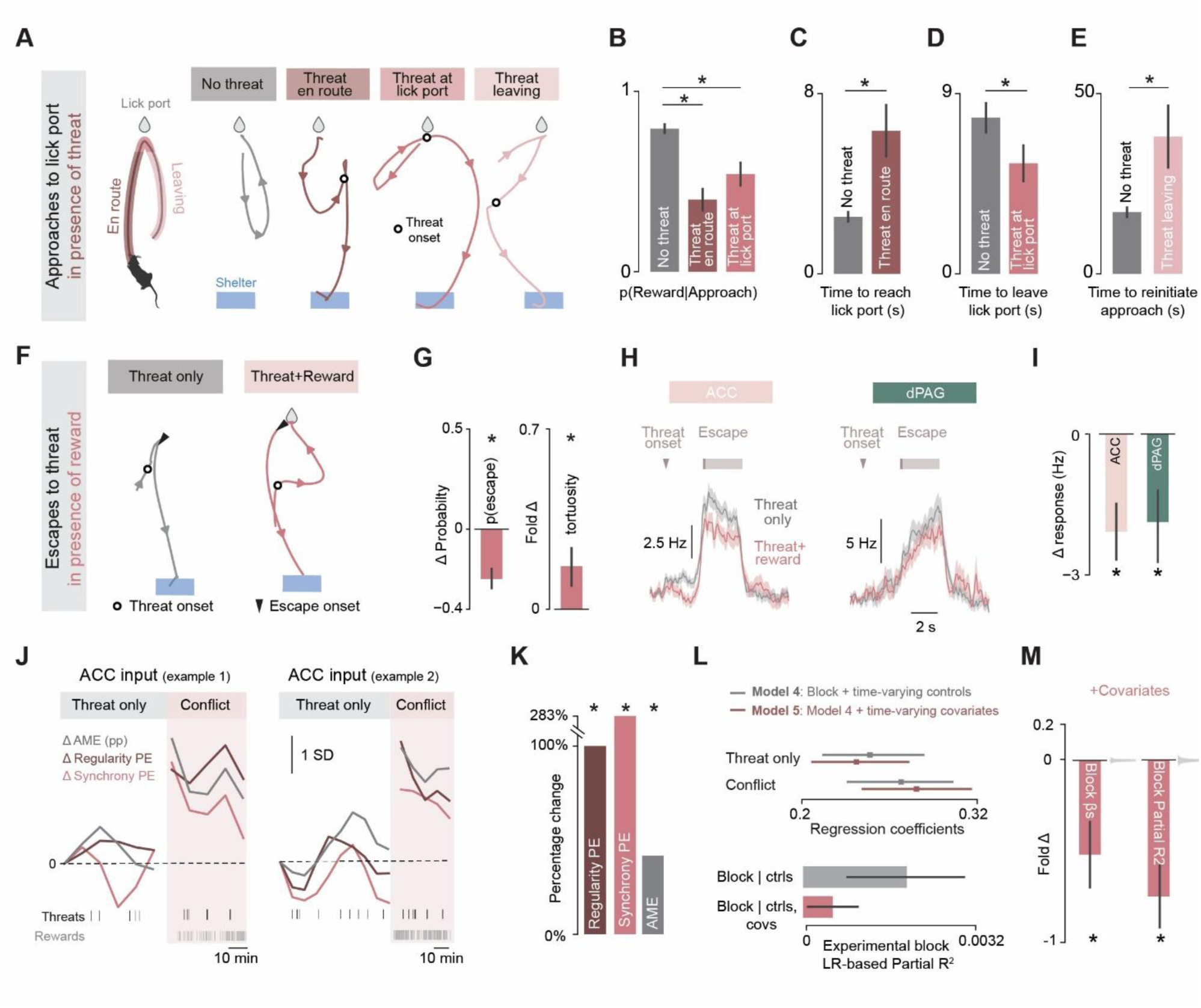
– Dynamic reweighting of ACC inputs onto dPAG during motivational conflict. **(A)** Example reward-seeking trajectories with no threat (grey) or with threat presented *en route* to the lick port (dark pink), at the lick port (pink), or just after leaving the lick port and before the next approach (light pink). The usual approach–reapproach loop (grey) is interrupted by threat in each case. **(B-E)** Session averages of the proportion of rewarded approaches (B) and trial averages of the time to reach (C), leave (D), and reinitiate (E) the next approach to the lick port, in the absence (grey) or presence (pinks) of threat. **(F)** Example escape trajectories in the absence (grey) or presence (pink) of the lick port. **(G)** Change in escape probability in response to visual looms (left) and fold change in escape-path tortuosity (right), in the presence vs. absence of the lick port **(H)** Average escape-aligned PSTHs of ACC (left) and dPAG (right) responses, in the absence (grey) or presence (pink) of reward. **(I)** Difference in escape response magnitude (presence − absence of reward) in ACC and dPAG. **(J)** Time-varying AME, regularity and synchrony postsynaptic effect for two example ACC inputs, across the threat-only and motivational-conflict blocks. **(K)** Percentage change between threat-only and motivational-conflict blocks in regularity postsynaptic effect, synchrony postsynaptic effect, and GLM-estimated AMEs of ACC inputs. **(L)** Regression coefficients of experimental-block contrasts (top) and likelihood-ratio-based partial R² of the experimental-block factor (bottom) in linear mixed models of AME magnitude, with random intercepts and either experimental block plus firing-rate controls (grey) or experimental block, controls, and temporal-statistics covariates (pink) as fixed effects. Error bars: bootstrap SEM. **(M)** Fold change in experimental-block coefficients (left) and partial R² (right) upon adding temporal-statistics covariates. Light grey kernel-density estimates: null distribution from shuffled covariates. Error bars: bootstrap SEM.

Reciprocally, the presence of reward shaped escape behaviour: escape probability decreased (Figure 6G, left; –25.2 ± 4.23 %; paired t-test, p < 0.0002) and trajectories were biased towards the lick port (Figure 6F) instead of the shelter-bound runs typical of unconflicted escape^6^, increasing escape tortuosity (Figure 6G; +23.6 ± 8.8 %; permutation test, p = 0.0038). This graded and bidirectional modulation of both behaviors suggests an active, ongoing weighting of safety against reward.

Single unit recordings during this assay confirmed that escape-related activity in both ACC and dPAG was modulated by the presence of reward (Figure 6I; change in average firing rate, ACC:-2.08 ± 0.595 Hz, paired t-test, p =1.26 x 10^−3^; dPAG: –1.87 ± 0.760 Hz, paired t-test, p = 0.0306), consistent with conflict-related signals reaching the defensive circuit. We then asked whether the functional impact of ACC inputs on dPAG also changed with behavioral context. Fitting separate GLMs to escape and conflict blocks revealed that ACC AMEs were substantially larger during conflict (Figure 6J-K; 0.0228 ± 0.00205 pp vs 0.0156 ± 0.00167 pp; paired t-test, p = 1.14 x 10^−5^) – that is, individual ACC spikes were stronger drivers of dPAG firing during motivational conflict.

This increase in functional impact was accompanied by an average increase in the postsynaptic effects of ACC synchrony and regularity by four– and two-fold, respectively (Figure 6K), both at a single-cell level (Figure 6J) and across the ACC population (by 0.00223 ± 0.000166 and 0.00388 ± 0.00125 for synchrony and regularity postsynaptic effect metrics respectively; paired t-test, p = 1.45 x 10^−36^ and 0.00205). Notably, this strengthening of functional drive occurred despite the modest reduction in mean ACC firing rate during escape, indicating that the increased influence of ACC reflected reorganisation of its temporal structure rather than a simple change in spike count. To test whether these temporal changes could quantitatively explain the elevated ACC AMEs during conflict, we adapted the LMM analysis used above (Figure 5L-N) by replacing brain region identity with block identity.

Both temporal statistic covariates received significantly positive coefficients (Supplementary Figure 13, permutation test, p < 0.001), and their inclusion substantially reduced the contribution of block identity to time-varying ACC AMEs: block contrast coefficients decreased by 43.9 ± 17.1 % (permutation test, p < 0.001; Figure 6L, top and 6M, left) and the overall explanatory value of block identity decreased by 70.8 ± 17.5 % (permutation test, p < 0.001; Figure 6L, bottom and 6M, right).

Together, these results show that the temporal-statistics framework established for inter-regional differences in functional connectivity also accounts for intra-regional changes across behavioural contexts: the same ACC inputs exert greater influence on dPAG during motivational conflict because they fire more synchronously and more irregularly under those conditions. Functional connectivity weight onto dPAG is therefore not a fixed property of each input pathway but a dynamic quantity, set on a moment-to-moment basis by the temporal organisation of presynaptic activity.

## Discussion

In this study, we asked how excitatory dPAG neurons integrate convergent inputs from five major afferent regions to compute flexible escape decisions. Combining simultaneous multi-region recordings during naturalistic behavior with subcellular synaptic mapping, dendritic biophysics, and biophysically constrained modelling, we found that functional connectivity onto dPAG is determined predominantly by the temporal statistics of presynaptic activity rather than synaptic placement or pathway identity. This single principle accounts for both the inter-regional hierarchy of input strength we observed across pathways — strongest for sensory midbrain (SC, IC), intermediate for cortical (ACC), and weakest for hypothalamic regions (VMH, PMd) — and the intra-regional, context-dependent reweighting of ACC influence on dPAG during motivational conflict. We propose that this principle emerges from the electrotonic compactness of dPAG neurons, which largely equalises the somatic impact of inputs across the dendritic tree and leaves temporal structure as the dominant determinant of input efficacy — providing a substrate for flexible, context-dependent input weighting on the timescales required for adaptive escape.

Reaching this conclusion required a multi-level approach integrating experimental and computational analyses across scales of the dPAG circuit. While GLMs are widely used to decompose task, stimulus and behavioural contributions to neuronal spiking, their application to integration hubs during ethological behaviour has remained limited. Here, we performed simultaneous high-density recordings from dPAG and each of its major input regions during naturalistic behaviour — yielding one of the largest existing datasets of dPAG single units and their afferents — and used biophysically constrained Bernoulli GLMs, anchored to experimentally measured dPAG membrane time constants, to estimate the functional influence of each input on dPAG spiking. The simultaneous recording of input and output regions was essential because it allowed each input’s influence to be conditioned on the activity of all other co-recorded populations, removing common-drive confounds that limit single-region or non-simultaneous approaches^75^. The resulting models recovered functional connectivity sufficient to reproduce threat-locked dPAG spiking without behavioural or sensory regressors, validating their use for quantitative comparison of long-range inputs in this system. Crucially, pairing these *in vivo* influence estimates with synapse-resolved anatomy, two-photon holographic dendritic stimulation and somato-dendritic whole-cell recordings, and biophysical modelling in the same circuit allowed us to interpret the statistical dependencies biologically: each level of analysis constrained the others, providing the resolution required to identify temporal dynamics as the dominant determinant of input influence.

The dominance of temporal dynamics over synapse location or pathway identity contrasts with the integration principles established in other well-studied regions, such as cortical and hippocampal neurons. In layer 5 pyramidal neurons, for example, distal inputs are substantially attenuated relative to proximal ones^63,64^, while clustered synapses on the same branch can trigger dendritic spikes that amplify their somatic impact^76,77^. In CA1 pyramidal neurons, the spatial segregation of entorhinal and CA3 inputs onto distinct dendritic compartments enables conjunctive computations that would not be possible if these pathways were intermingled^78^. In contrast, dPAG neurons appear to operate by a fundamentally different logic: somatic responses are largely invariant to synapse location, and synapses from different input regions, despite organisational biases, exert influence in proportion to the temporal structure of their activity rather than their dendritic position. We propose that this regime arises from the electrotonic compactness of dPAG neurons – a consequence of high input resistance, long effective length constants, and sparse, weakly branched dendrites – which together cause these cells to behave closer to single electrical compartments than spatially extended cables. The reproduction of all measured biophysical properties by passive cable models suggests that active dendritic conductances, if present, are either modest or uniformly distributed^79,80^. More generally, this finding also reveals how relatively unspecialised, isodendritic morphologies^81^ can support their function: rather than relying on distance-dependent filtering or branch-specific computations that require precise axonal targeting, the dendritic tree provides broadly uniform coupling across the arbor, implementing a mechanistically simple form of ‘synaptic democracy’ in which presynaptic temporal dynamics dominate spiking output. This result further extends previous work demonstrating that SC-dPAG synapses are weak and unreliable and require recurrent amplification within SC to generate sufficient synchronous drive^4^. Our multi-pathway comparison suggests that this synaptic weakness may be a general property of dPAG inputs: if all afferent synapses are individually weak, then temporal coincidence becomes the principal route to driving dPAG firing, and the network-level dynamics of each input region — rather than the strength of any single synapse — determine its functional weight.

Given that all inputs have comparable access to the soma, the variable that differentiates them is the temporal structure of their activity. Synaptic currents summate when they overlap in time, and the degree of overlap depends on both within-neuron firing patterns and across-neuron coordination — properties that we found to differ systematically across dPAG afferents. SC and IC neurons fire with high spike irregularity, producing an excess of short interspike intervals that favor temporal summation of successive spikes from the same neuron, and with high population synchrony, causing spikes from different neurons to arrive within the postsynaptic integration window. The synchrony of SC and IC is itself consistent with the recurrent connectivity of SC^82,83^ and with shared feedforward drive producing localised synchrony in IC^84^. VMH and PMd, by contrast, fire closer to Poisson processes with low synchrony, producing temporally dispersed inputs that summate poorly. These differences in temporal statistics accounted for ∼64% of the variance in functional connectivity, indicating that input strength arises not as a fixed property of any pathway but as an emergent consequence of how presynaptic populations interact with the integration constraints of dPAG neurons.

The temporal sensitivity we describe in dPAG resembles coincidence detection in other systems, but differs in a critical respect: it seems agnostic to pathway identity. For example, neurons in the medial superior olive compare two specific input streams to detect microsecond interaural disparities^85,86^, and in the SC, the response of multisensory neurons depends on the cross-modal nature of the coincidence^87,88^. In these cases, the identity of the converging streams is central to the computation. In contrast, dPAG neurons integrate temporal coincidences equally for all streams: any pathway can drive spiking strongly if its activity is sufficiently synchronous, and the spatial configuration of synapses contributes minimally. This input-agnostic regime implements a form of weighted summation across pathways without requiring pathway-specific synaptic mechanisms — coincident activity from a single input is, in principle, equivalent to coincident activity distributed across several.

A direct implication of this principle is that the inter-regional hierarchy reported here is not a fixed feature of the circuit, but a snapshot under one set of behavioural conditions. Because functional weights are set by presynaptic temporal dynamics rather than by structural features of the connections themselves, the hierarchy should reorganise whenever those dynamics shift. For example, while hypothalamic inputs were weak in our conditions, in states that synchronise or burst-coordinate their activity — such as sustained live predator presence or specific neuromodulatory regimes^22,36^ — the same connections could in principle dominate dPAG output. The escape circuit appears not to assign privileged status to any single pathway: any input region can become a strong driver of dPAG firing.

The combination of input-agnostic temporal integration and contextually variable presynaptic dynamics has a direct implication the computation of escape decisions: dPAG can rapidly reweight its inputs without modifying any synaptic connection. We observed this directly in our motivational-conflict experiments. When mice had to weigh reward against threat, individual ACC spikes became more effective at driving dPAG firing than during pure escape. The change was accompanied by an increase in the synchrony and firing irregularity of ACC neurons, and most of the increased ACC influence on dPAG could be explained by these temporal features alone. The reweighting therefore arose from reorganisation of presynaptic temporal structure rather than from changes in the synapses themselves. This architecture provides a substrate for the rapid, reversible modulation of escape that animals require: factors known to alter escape probability — hunger, stress, prior experience^1,8^, and acute threat appraisal — can act by shaping the temporal dynamics of dPAG afferents, rewiring functional weights on a moment-to-moment basis without engaging plasticity at the dPAG synapses themselves.

More broadly, our findings illustrate how subcellular biophysics can be tuned to shape the role a neuron plays in its circuit. The electrotonic compactness that makes dPAG neurons input-agnostic and temporally driven is a substrate for the specific computation they perform — flexible, multi-source integration of competing signals into a single survival decision.

Convergence hubs throughout the brain may operate by similar logic, with the integrative properties of single neurons calibrated to the demands of distributed input streams that must be combined, weighted, and reweighted on behavioural timescales.

## Limitations

Several caveats merit explicit consideration. First, the functional connectivity estimates we report are statistical inferences from joint spiking patterns rather than direct measurements of synaptic transmission. Although simulations established that input temporal statistics causally shape GLM-estimated coupling, the framework would benefit from complementary *in vitro* measurements of pathway-specific synaptic strength. Differences in synaptic properties between input pathways could contribute to the residual variance not captured by our temporal-statistics covariates, and resolving this would require systematic comparison of evoked synaptic responses across pathways. Such experiments are, however, technically challenging, requiring single synapse stimulation, ideally through opsins expressed uniformly across pathways to ensure that evoked responses can be quantitatively compared. Second, our conclusions about dendritic integration of synaptic input in dPAG rest predominantly on biophysical modelling rather than on direct experimental measurement. Techniques such as focal electrical stimulation or two-photon glutamate uncaging would be needed to test location-dependent synaptic integration experimentally; dendritic patch-clamp at more distal locations to measure attenuation over larger distances; and pharmacological blockade to rule out contributions from voltage-gated dendritic conductances. Our passive compartmental models reproduce all measured properties without invoking active mechanisms, but small non-uniform contributions from active conductances cannot be ruled out by our current data. Third, we focused exclusively on excitatory inputs and excitatory dPAG neurons, leaving the role of inhibition unaddressed. Long-range inhibitory projections to dPAG, as well as excitatory inputs onto local inhibitory dPAG neurons, are likely to shape the integration of the long-range excitatory drive we characterised here, and incorporating these pathways will be an important direction for future work. Finally, although our motivational-conflict experiments demonstrate that ACC functional connectivity reweights with behavioural context on rapid timescales, broader sampling of internal states will be needed to test the full scope of contextual reweighting predicted by the framework.

## Methods

### Mice

All experiments were carried out under the UK Animals (Scientific Procedures) Act of 1986 (PPLs: 70/7652 and PFE9BCE9), following local ethical approval by the Sainsbury Wellcome Centre Animal Welfare Ethical Review Body.

Mice of the vGluT2-ires-Cre driver line (Jackson Laboratory, Stock No. 016963), and mice from crosses of vGluT2-ires-Cre with either the R26 EYFP (Jackson Laboratory, Stock No. 006148) or Ai14, (Jackson Laboratory, Stock No. 007914) reporter lines were used in this study (both male and female mice, age 5–12 weeks. All mice were bred in-house in the Sainsbury Wellcome Centre Neurobiological Resource Facility. Mice had free access to food and water on a 12:12 h light:dark cycle (ambient temperature 24 °C, humidity 47 relative humidity) and were tested during the dark phase.

### Surgical procedures

All surgical procedures, including chronic implantation of devices and viral injections were performed as described in^31^. Briefly, general anaesthesia was induced and maintained with isoflourane (0.5-4%). Carprofen (5mg/kg) was administered subcutaneously before surgery for analgesia. During surgery mouse was secured in a stereotaxic frame (Kopf Instruments, model 1900 or 963). Viral constructs were loaded in a pulled glass pipette (10 μl Wiretrol II) and injected using a hydraulic micromanipulator coupled to an injection system (Narishige, MO-10). Implants were affixed using light-cured dental cement (3M, RelyX Unicem 2) and the surgical wound closed with surgical glue (Vetbond). Mice were placed in a clean, heated recovery cage after surgery, and returned to their home cage after full recovery from anaesthesia.

### Viruses

The following viruses were used in this study and are referred to by contractions in the text. Optotagging experiments AAV2/2-EF1a-DIO-hChR2(H134R)-EYFP (3.6×10^12 VG/ml, UNC Gene Therapy Vector Core), AAVrg-CAG-Flex-Flpo (4.5 × 10^14 VG/ml, Sainsbury Wellcome Centre Viral Vector Core), AAV2/1-EF1a-fDIO-ChrimsonR-tdTomato (5.8 × 10^13 VG/ml, Sainsbury Wellcome Centre Viral Vector Core) and AAV2/1-CamKII-Cre (1 × 10^13 VG/ml, Addgene); mapping of spatial organisation of synapses: AAV2/2-EF1a-DIO-synatophysin-mCherry (2.48 × 10^13^ GC/ml, Sainsbury Wellcome Centre Viral Vector Core), AAV2/1-CAG-Flex-Flp (5 × 10^12 GC/ml, Sainsbury Wellcome Centre Viral Vector Core) and AAV2/1-TRE-fDIO-GFP-IRES-tTA (5.80 × 10^13 GC/ml, Sainsbury Wellcome Centre Viral Vector Core, from Addgene plasmid #118026); holographic stimulation experiments: AAV2/2-CAG-Flex-Flp (diluted down to 10^10^ GC/ml, Sainsbury Wellcome Centre Viral Vector Core) and AAV2/1-EF1a-fDIO-hChR2(H134R)-EYFP (9.2 × 10^14^ GC/ml, Sainsbury Wellcome Centre Viral Vector Core)

### Single-unit recordings in freely moving mice

#### Surgery and histology

vGluT2-Cre mice were injected with AAV-EF1a-DIO-hChR2-EYFP in dPAG to allow for optotagging of glutamatergic dPAG neurons. For optotagging of glutamatergic presynaptic cells, AAVrg-CAG-Flex-Flpo was also injected in dPAG, and AAV-EF1a-fDIO-ChrimsonR-tdTomato injected in SC, IC, VMH and PMd. 10:1 mix of AAV-EF1a-fDIO-ChrimsonR-tdTomato and AAV2/1-CamKII-Cre was injected in ACC. At least 8 weeks was given for virus expression before a second surgery during which the optic fibre and Neuropixels probes were implanted. A 2.5mm long, 400um diameter optic fibre of 0.5 NA (Newdoon) was implanted unilaterally above the dPAG with a mediolateral angle of 35° and entering the brain from the contralateral side. One 4-shank Neuropixels2.0 probe was chronically implanted in PAG, SC, IC and PMd with a 40° anterior-posterior angle, and another either single-shank or 4-shank Neuropixels2.0 probe implanted in ACC and VMH with a –35° anterior-posterior angle. Probe shanks were coated with DiD (1mM in ethanol, Invitrogen) prior to insertion, to allow for track identification.

At the end of the experiment, the mouse was perfused and the brain was imaged using serial micro-optical sectioning 2-photon tomography^89^ to confirm the location of probe implantation and expression of ChR2 and Chrimson.

#### Behavioural procedures

All behavioural experiments were performed in an elevated white, circular Perspex arena (92 cm diameter) with a red see-through Perspex shelter (20 x 10 x 10 cm; 12 cm wide entrance) placed at the edge of the arena and low luminance conditions (typically 1.2 lux)^6,31^. Video recording (40 fps, acA1300-60gmNIR, Basler), sensory stimulation and reward delivery were controlled with Bonsai^90^. Mouse position was tracked online using Bonsai centroid tracking.

Escape assay was performed as previously described^6,31^. The mouse was introduced in the experimental arena and, before presenting any threatening stimulus, it was left exploring for a 7 min habituation period, during which it had to enter the shelter at least once. During a single session, multiple stimuli were delivered with an inter-stimulus interval of at least 180s. The visual threat stimulus was a sequence of 5 expanding dark circles that expanded linearly over 400 ms then maintained the same size for 250 ms and repeated with a 500 ms interstimulus interval. The visual stimulus was presented on both an overhead projector screen and a side computer monitor. The auditory threat stimulus was either a 9s long 10kHz tone or a sequence of 3 10kHz tones whose amplitude increased linearly over 1s and repeated with a 500 ms interstimulus interval (sound intensity 80db^31^). A standard computer speaker positioned next to the arena was used to deliver non-threatening, naturalistic auditory stimuli, which were .wav files played via the sound card of the acquisition computer and fed through an amplifier.

Air-puff ports were positioned at the edge of the arena and connected to compressed air supply via a TTL-controlled valve that controlled the timing and duration of air-puffs.

A small peristaltic pump (WPM1-P1AB-WP, Welco) was used for condensed milk delivery at a nose-poke lick-port that was positioned at the edge of the arena only during the reward block. Condensed milk was placed in a small beaker sealed with parafilm below the arena. The position of the mouse was tracked online, and reward delivery was contingent on the mouse first moving out of a predefined region near the lick port (typically 40 cm x 40 cm) before returning to perform a nose poke at the port. Food-restricted mice were trained to obtain milk from the lick port in this manner over 2-3 60 min long sessions one week prior to probe implantation.

To elicit hunting behaviour, live adult black field crickets (Gryllus sp., Blades Biological) were placed on the arena floor (typically 3-4 crickets per session).

A female, adult C57BL/6J mouse was placed in an enclosed white Perspex box that was positioned next to the arena and had small holes along the walls, such that the recorded mouse could smell, hear and see the female mouse, but not physically interact with the female mouse in any way.

Filter paper loaded with 10ul of 10% 2,4,5-trimethyl thiazoline (TMT, Sigma) diluted in water was placed in a small petri dish with holes in its lid. Unloaded filter paper was placed in another similar petri dish, and both were placed on the floor of the arena.

#### Data Acquisition

Neuropixels probe recording and optotagging were performed as described in^31^Briefly. signals from the 2 Neuropixels 2.0 probes were recorded using SpikeGLX (https://billkarsh.github.io/SpikeGLX/) at a sampling rate of 30 kHz and high pass filtering at 300 Hz. A rotary joint (either AHRJ-OE_PT_AH_12_HDMI or AHRJ-OE_1×1_PT_FC_24_HDMI+4, Doric) was used to prevent twisting and torsion cables. Active recording sites were chosen based on expected probe depths and a whole probe activity survey done prior to recording sessions. For optotagging, the following stimulation protocol was used: 10ms long, 639nm wavelength pulse (Vortran Stradus VersaLase); 10ms long 472 nm wavelength pulse (Vortran Stradus VersaLase); 10ms long 472 nm wavelength pulse with preceded by a 250ms long 693nm wavelength pulse^91^. Pulse types were repeated 40 times in a randomized order and further repeated at 4 different laser intensities (powers at the fibre optic tip: minimum 0.75 mW, maximum 20mW for 693nm; minimum 0.3 mW, maximum 30mW for 472 nm).

#### Analysis of behavioural data

Escape success and failure and escape onset were manually annotated based on the animal making a full escape to the shelter. Escape was defined as a continuous movement consisting of a head orientation followed by full-body turn and run towards the shelter^4,6^. Escape onset was defined as the first video frame in which the mouse makes the initial head orient^4,6^. Escape probability was calculated as the proportion of trials in a session where stimulus triggered escape to the shelter. For each escape, the following metrics were quantified: reaction time, defined as the latency between stimulus onset and escape onset; maximum escape speed, defined as the peak speed reached between escape onset and shelter entry; and path tortuosity, defined as the ratio of the path length taken to the shortest distance to the shelter

Overall engagement in reward-seeking was quantified as 1) the rate of reward, based on the number of rewards divided by the duration of time the lick port was in the arena, and 2) the proportion of time spent near the lick port, based on mouse location from the tracking data. Metrics were quantified for each session and averaged across sessions and mice. Time spent in the same ROI prior to lick port addition with only the shelter in the arena (i.e. the ‘threat only’ block) was used as the baseline for comparison using a paired t-test.

Approaches to the lick port during the reward period were detected from tracking data as continuous periods (> 700 ms) during which the distance of the mouse to the lick port decreased quickly (> 30 cm/s) and with the trajectory pointing towards the lick port (< ± 1 rad). Approach trials that were closest in time to a reward delivery event and within 8 s of the reward delivery event were defined as rewarded approaches i.e. approaches that led to reward delivery. Approach trials that were closest in time to a threat presentation and within 20 s of the event were defined was approaches in the presence of threat. For each approach, the following metrics were quantified: the time between approach onset and lick port arrival (“time to reach lick port”), the time spent at the lick port from lick port arrival to leaving (“time to leave lickport”), and the time from leaving the lick port to the start of the next approach (“time to reinitiate approach). The probability of obtaining a reward given an approach was quantified as the fraction of rewarded approaches in each session.

Periods of cricket interaction were manually annotated from the behavioural video and classified into 1) detection, the period between cricket entry in the arena and first contact between the mouse and cricket, 2) capture, the period during which the mouse chases the cricket and secures possession of the cricket, which often involves bringing the cricket back into the shelter 3) immobilisation, during which the mouse begins to tear apart the cricket, with small bouts of chasing occurring largely within the shelter as the cricket escapes from the mouse’s grasp 4) consumption, during which the cricket is no longer moving, and the mouse is eating the cricket. Proportion of time spent in each type of cricket interaction were quantified for each session and averaged across sessions and mice.

#### Isolation and anatomical registration of single units

Neuropixels data were preprocessed using CatGT (https://github.com/billkarsh/CatGT) as such: band-pass filtering between 300 Hz and 5000 Hz with an order-12 biquad filter, local common average referencing on an annular area of 300um outer radius and excluding an inner radius of 100um, and zeroing-out large amplitude excursions that occur across most channels. Spike sorting was done using Kilosort 2.5^92^ and manually curated using Janelia Rocket Cluster 3.0^93^. The brain was registered to the Allen Brain Atlas using brainreg^94^ and tracks manually reconstructed in brainreg-segment. Active sites were assigned brain regions based on the registration and additionally adjusted based on electrophysiological signatures.

#### Identification of opto-tagged units

Units were classified as optotagged as previously described^31^. Briefly, Putative VGluT2+ cells had to spike within 5 ms of blue laser stimulus end in at least 50% more trials than before the stimulus (i.e., light-triggered spikes happen in at least 50% more trials above the base occurrence of spontaneous spikes) and with a maximum jitter (IQR) of 3 ms. Putative dPAG-projecting VGluT2+ cells had to spik within 10ms of red laser stimulus end in at least 40% more trials and with a maximum jitter of 4ms.

#### Firing rate modulation of units

To identify units that showed rate modulation to the various behavioural conditions, firing rates of each unit during specific behaviours were quantified in 1 s bins and compared using Bonferonni-corrected paired t-tests. Only time periods where the mouse was outside the shelter were used. Escape-modulated units were defined as units whose firing rate within 5s of threat presentation and 3s of escape onsets were significantly different from during exploration bouts, within the ‘threat only’ block. Reward-modulated units were defined by comparing firing rates when the mouse was near versus far from the lickport during the ‘reward’ block; cricket-modulated units by comparing firing rates when the mouse was engaged in cricket hunting versus non-cricket related behaviour during the ‘hunting’ block; female-modulated units by comparing when the mouse was near versus far from the female during the ‘female’ block; TMT-modulated units by comparing when the mouse was near the TMT-loaded petri dish versus just before the ‘TMT’ block; and sound-modulated units by comparing during naturalistic sounds to just before the ‘sounds’ block. To test whether the different behaviours produced distinct distributions of modulation across regions, the proportion of modulated cells in each region was quantified and normalised within each behaviour to obtain modulation compositions. The sum of squared Euclidean distances between behaviours was used as the test statistic, and significance assessed using a permutation-based test in which the behaviour labels of each cell were randomly shuffled.

#### Peristimulus trial-averaged responses

Spike trains were aligned to trial starts, binned at 10ms and smoothed with a 30ms causal gaussian filter. The trial-wise firing rates were baseline-subtracted with the average firing rate in the 1s baseline before each trial start and then averaged across trials to obtain the peristimulus trial-averaged histogram (PSTH). For escape trials, the spike trains aligned to threat onset were additionally resampled in time in order to align escape onset and shelter entry across trials (“time-warped PSTHs”^25^). For visualisation, PSTHs were normalised to their maximum absolute value (i.e. maximum or –minimum, whichever is larger), to account for variability in firing rate changes across cells. For identification of threat– and escape-responsive units, PSTHs of all units across brain regions and recording sessions were reduced to the top 200 principal components and grouped into non-responsive, inhibited and excited using k-means clustering on specific time periods. Threat-responsive units showed short-latency excitation within 500 ms of stimulus onset, and escape-responsive units showed excitation within the first one-third of the escape run.

#### Comparisons of neural response magnitudes during threatening stimulus presentation

Comparisons of escape response magnitudes in the presence versus absence of rewards were done on neurons with escape responses, with mean responses computed on the time-warped PSTHs over the full escape duration.

#### Quantification of intra-neuron temporal statistics

Each neuron’s interspike intervals (ISIs) were extracted as the time between consecutive spikes, and normalized by the mean ISI. Each neuron’s ISI distribution was obtained by quantifying either the proportion of raw ISIs in every 1 ms bin up to 100 ms, with an overflow bin for ISIs above 100 ms, or the proportion of normalized ISIs in bins up to 1, with an overflow bin for ISIs above the mean ISI. ISI distributions were averaged across neurons in each brain region for visualization. ISIs were also modelled as arising from a shifted gamma renewal process, with a shape parameter α, rate parameter 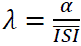 and a refractory period τ_refr_, such that the refractory-corrected ISIs satisfy 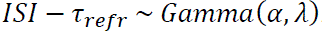. Parameters were estimated by fitting the normalized ISIs with a shifted gamma distribution.

To calculate the regularity postsynaptic effect, The average peak of the postsynaptic response for a presynaptic spike train was estimated by integrating dPAG’s temporal summation curve (obtained from NEURON^95^ simulations of inputs at the soma) against the distribution of raw ISIs, and normalized by the integration against a Gamma distribution of α = 1 and λ = mean firing rate of that spike train as a measure of the effect of the ISI distribution shape on postsynaptic response.

#### Quantification of inter-neuron temporal statistics

The cross-covariance between pairs of simultaneously recorded neurons in the same brain region was evaluated at 0 ms time lag, to measure the absolute rate of coincidence of spikes that is above chance from the mean firing rates, expressed here per second instead of per 1 ms time bin. For each neuron, pairwise cross-covariances with other neurons were averaged, to obtain a measure of the average above-chance rate of coincidence of its spikes with other neurons.

To estimate the effect of population synchrony on postsynaptic response size while taking higher-order interactions (i.e. beyond pairs of neurons) into consideration (synchrony postsynaptic effect), the proportion of each neuron’s spikes that occurred within 0-100 ms of another spike from any neuron in a subpopulation of the region was quantified to obtain equivalent ISI distributions for each neuron that are now in the context of the firing of other neurons in the same region. Each subpopulation was randomly sampled from neurons simultaneously recorded in the same region, from subpopulation sizes of 1 (i.e. with only the neuron’s own spikes) to all the simultaneously recorded neurons in the same region, with up to a maximum of 1000 unique combinations for each subpopulation size. The ISI distributions for that neuron at each subpopulation size were then averaged across the randomly sampled combinations, resulting in an average ISI distribution at each subpopulation size for each neuron. The dPAG temporal summation curve was then integrated against each average ISI distribution to obtain the average expected peak response size for that neuron at each subpopulation size. The rate of increase (k) of expected response (y) with population size (n) was estimated by fitting a shifted saturation curve with the equation *y* = *y*_0_ + (*y*_*max*_ − *y*_0_)(1 − *e*^−*k* × *n*^), where y_max_ is the peak of the temporal summation curve at 0 ms (∼1.948).

Population saturation rate constants were also recomputed using circularly-shifted spike trains that preserved auto-correlation structures and firing rates within each neuron, while disrupting millisecond-precise spike alignment and co-modulations of firing rates across neurons. All spikes belonging to each neuron were shifted in time relative to the spikes of other neurons by a constant, random lag of at least 80 s. Shifted spike times that exceeded the recording duration were circularly shifted to the start of the recording.

#### Modelling of dPAG spiking probability from input activity using GLMs

A generalised linear model with a logistic link function was used to predict the probability of spiking in dPAG neurons given simultaneously recorded spike trains of neurons from known input regions to dPAG as predictors. Spike trains used were ongoing spike trains from the whole recording session, with 50% of threat presentation trials excluded for use as the test set, excluding the optotagging block and extended periods in which the mouse remained in the shelter. Each binary spike train predictor was convolved with an exponential decay filter with a decay time constant of 18 ms, matching the average membrane time constant of vGluT2+ dPAG neurons estimated *in vitro*. L1 regularisation was put in place to promote sparsity. Synaptic delay and time to peak of synaptic current was modelled by time-shifting the presynaptic spike trains relative to the PAG spike train by 3 ms.

Model prediction accuracy was assessed by computing the Pearson correlation coefficient between trial-averaged predictions of spike probability and real PSTHs (25ms causal gaussian filter), aligned to threat and/or escape onsets depending on what the dPAG neuron was responsive to. Significantly predicted cells for which the model successfully captured neural modulations above chance were defined as cells for which the Pearson’s correlation coefficient was significantly larger than the null distribution obtained from shuffling the input matrix, after Bonferroni correction.

#### Statistical analysis of regional differences

Model coefficients (β) were converted to average marginal effects (AME) to express the effect of each predictor on the probability scale. For each predictor, the marginal effect at every time bin was computed as the derivative of the predicted probability with respect to the predictor (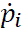), averaged across all time bins and multiplied by 100 to obtain the AME in percentage points, representing the average percentage point change in spike probability per unit change in the predictor:

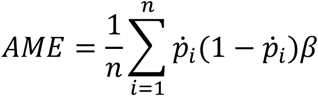

A cut-off of 3 x 10^−4^ percentage-point AME was used to define excitatory functional connections (see figure S4C).

Regional differences in the proportion of excitatory functional connections were assessed by comparing each region’s proportion to the overall average, using a permutation-based test in which subsets of neurons, matched to the number of recorded cells in each region, were randomly sampled without replacement from the full set of recorded neurons regardless of region identity.

Regional differences in the magnitudes of excitatory functional connections were assessed using a wild bootstrap ANOVA and post-hoc wild bootstrap tests of mean differences, with two-stage false discovery rate correction for multiple comparisons. In each wild bootstrap replicate, residuals relative to the pooled mean (i.e., under the null hypothesis of equal means) were resampled with replacement within each region and multiplied by Rademacher weights (±1 with equal probability), preserving the variance structure for each region. The Rademacher-weighted residuals were then added back to the pooled mean to generate resamples, and the F statistic or difference in group means was calculated for each bootstrap replicate to form the empirical null distribution.

To test whether AMEs of the optotagged inputs and AMEs to optotagged PAG cells originated from the same underlying distribution as those of untagged cells, permutation-based tests were used to assess exchangeability (i.e. null hypothesis of equal mean, variance and overall distributional shape). Permutation-based proportion and Kolmogorov–Smirnov tests were used to compare proportions of excitatory functional connections and the distribution of their magnitudes respectively.

#### Linear mixed-effects model

A linear mixed-effects model was used to estimate the contribution of multiple fixed effects to AME magnitude while modelling non-independence of AMEs coming from the same GLM (“groups” of AMEs) using random intercepts for each group. The random effect captures potential differences across GLMs that are not easily quantified but can potentially influence AME magnitudes, such as the proportion of the PAG’s true presynaptic cells that had not been simultaneously recorded (i.e. proportion of ‘missing’ predictors) or proportion of unconnected cells that were simultaneously recorded and partially correlated with the true presynaptic cells (‘nonsense’ predictors), both of which could bring down AME magnitudes assigned to recorded functionally connected input cells. The fixed effects included brain region, input firing rate on a log-scale and PAG firing rate on a log-scale (‘control’ fixed effects), and an index for the postsynaptic effect of enrichment of short inter-spike intervals and an index for the postsynaptic effect of intra-region neuronal synchrony (‘covariate’ fixed effects), depending on model specification.

Significance of the fixed effects were assessed using a likelihood ratio test, comparing the log-likelihood of the full model to that in the nested, reduced model in which the fixed effect of interest was removed. The resulting likelihood ratio was also converted to a Cox and Snell pseudo-R², which ranges from 0 to less than 1, to provide a measure of the incremental improvement in model fit attributable to inclusion of the fixed effect. This was calculated as:

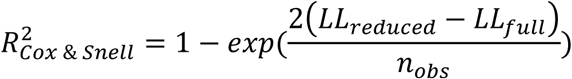

Standard errors of the likelihood ratio-based pseudo-R² and model coefficients were estimated using the nonparametric bootstrap with resampling stratified by groups.

Changes in the contribution of the brain region factor in the presence of the control or covariate fixed effects were quantified as the fold difference in brain region-associated likelihood ratio-based pseudo-R² and average fold difference in brain region-related model coefficients. Significance was assessed with permutation tests in which within-group factors (i.e. factors that vary within each group) were independently permuted within each group, and group-level factors (i.e. factors that are constant within each group, such as PAG firing rate) were shuffled across groups.

Partial residual plots were constructed by computing the partial residuals for a given predictor of interest X_i_ as 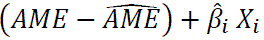 and plotting the moving mean (over a sliding window of 250 observations) against X_i_. The partial residuals were also additionally shifted by the LMM intercept and contribution of all other covariates at their mean value, for alignment with the plot of 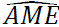 against X_i_.

#### Analysis of variation in ACC-dPAG functional connectivity over time

Separate GLMs were additionally trained on spike trains during the escape block and motivational conflict block, to compare differences in ACC-dPAG functional connectivity across the two behavioural blocks. Successive 10-minute windows were used to capture the evolution of AMEs over time. Properties of the GLMs were otherwise identical to the full-session GLM above. Only dPAG units whose firing had been significantly predicted above were used. A linear mixed-effects model with modified random and fixed effects was used to capture the variation in AMEs over time for each ACC-dPAG pair. Random intercepts were now assigned to each input of each GLM to absorb time-stable, across-pair differences in AMEs. Fixed effects included experimental block identity, and the same ‘control’ and ‘covariate’ fixed effects as above, but calculated for each 10-minute window and mean-subtracted for each input to capture dynamic fluctuations over time independently of stable differences across ACC units.

#### Simulations of feedforward leaky-integrate-and-fire neural networks

Simulated spike trains were generated from simulating a feedforward network composed of conductance-based leaky integrate-and-fire neurons in Brian 2^96^, in which presynaptic neuron populations with specified spiking statistics were synaptically connected to postsynaptic neurons with specified connection probabilities and synaptic conductances. Spiking in the postsynaptic neurons was driven solely by passive integration of inputs from these defined presynaptic neuron populations pushing the membrane potential beyond a predefined spike threshold. Properties of the postsynaptic, leaky integrate-and-fire PAG neuron were experimentally derived wherever possible, and set as the following unless specified:

|  |  |
| --- | --- |
| Membrane capacitance | 0.0514 nF |
| Leak conductance | 2.84 nS |
| Threshold membrane potential | -38.9 mV |
| Reset membrane potential | -64.5 mV |
| Resting membrane potential | -75 mV |
| Refractory period | 2 ms |

Synapses were modelled as exponential synapses, with the synaptic conductance showing an instantaneous rise and exponential decay of a time constant of 2 ms. Peak conductance was set to 0.5 nS unless specified. Firing in presynaptic neurons was modelled as Poisson processes unless specified, and with a rate of 5 Hz unless specified. The number of connected presynaptic neurons was adjusted to maintain postsynaptic firing rate within 10-15Hz, to match experimentally observed PAG firing rates. For non-Poisson spiking, spike trains with varying interspike-interval statistics were generated by randomly sampling interspike intervals from a Gamma distribution whose shape parameter α was varied (rate parameter λ = α x firing rate), with a burn-in duration of 100 s. Spike trains with varying synchrony were generated by sampling from dichotomized Gaussian models with specified correlation coefficients and firing rates per neuron^97^.

### Mapping of spatial organisation of synapses

#### Mice and viruses

Viral labelling was done in vGluT2-Cre mice to label glutamatergic synaptic terminals onto glutamatergic dPAG cells. To label presynaptic terminals, AAV-EF1a-DIO-synatophysin-mCherry was injected into an input region of interest (SC, IC, VMH, PMd or ACC). Sparse, bright labelling of dPAG was achieved by injecting AAV-CAG-Flex-Flp injected in an input region of interest, from which it inefficiently jumps trans-synaptically in the anterograde direction to PAG, and AAV2/1-TRE-fDIO-GFP-IRES-tTA injected in dPAG.

#### Brain tissue clearing using CUBIC

Mice were transcardially perfused with 4% paraformaldehyde 2 weeks after viral injection, and brains were extracted and post-fixed overnight at 4°C. Brains were sectioned coronally on a vibratome to obtain 150 µm sections of the input region of interest, which were then mounted on glass slides and imaged with an epifluorescence microscope (Zeiss Axio Imager 2) to confirm accurate targeting of the synaptophysin virus. In addition, a thick 1.7mm ‘slab’ was taken at the level of PAG to preserve continuity of neurites traversing in the anterior-posterior direction, and optically cleared using the CUBIC protocol^53^ to facilitate imaging of the entire PAG as a single, continuous z-stack. In brief, the sample was incubated in 0.5x CUBIC L (diluted 1:1 in PBS) for 8 hours at room temperature, followed by 1 x CUBIC L at 37 degrees for 5 days, to achieve delipidation. The sample was then given 3 x 10-minute washes in PBS and further stored in PBS at room temperature. 3 days before imaging, the sample was transferred to 0.5 x CUBIC R (diluted 1:1 in dH2O) for 8 hours, glued down onto a petri dish and incubated in CUBIC R for 2 days at room temperature to achieve refractive index matching. The CUBIC R solution was slightly diluted with dH2O until it reached a refractive index of 1.457, to stay within range of refractive index mismatch correction of the objective lens.

#### Confocal imaging

A Leica SP8 confocal microscope equipped with the Leica HC FLUOTAR L 25X/1.00 IMM (NE=1.457) MOTCORR VISIR objective optimized for CLARITY-treated specimens (Leica, article no. 507703) was used to obtain high-resolutions images of the CUBIC-cleared PAG sample. EGFP and mCherry were imaged simultaneously using 488 nm and 587 nm from a white light laser for excitation, and emission collected at 493-571nm or 592-789nm respectively. Laser power was adjusted across Z planes to maintain fluorescence intensity with depth while avoiding oversaturation and channel bleed-through. The sample was imaged at 0.216 um resolution in X and Y, and 2 um in the Z axis, with a field of view covering dPAG and neighbouring areas to contain all visible dendrites, over typically 40 hours of total imaging time.

#### Reconstructions

Confocal image stacks were stitched, merged and denoised. Neurons were then traced in 3D from the EGFP channel in a semi-automated manner (Vaa3D with the TeraFly plug-in for large datasets^54^). Dendrite radius was estimated automatically based on signal intensity and manually proofread. Colocalization of signal in the EGFP channel and the mCherry channel (containing potential presynaptic boutons) occurring along the neuron reconstructions was automatically detected using the Synapse Detector plug-in in Vaa3D^98^, and then manually curated. Morphometric analysis of the resultant neuron reconstructions and synapse locations was done in Python, using the NeuroM package and custom code.

#### Statistical analysis

For each reconstructed dPAG neuron, the path distance from the soma to each putative synapse was quantified. For each input region, a bootstrap-based Kolmogorov-Smirnov test was used to compare the average distribution of synapse path distances from the soma against the average distribution of path distances in the dendritic tree, to test if it was equally likely for a synapse to occur at every path distance from the soma. The observed Kolmogorov-Smirnov statistic was compared against a null distribution obtained by sampling locations with replacement from each cell’s dendritic tree at an equal probability (i.e., under the null of independent and uniform, random placement along the dendrite) and recalculating the statistic over 10,000 resamples. Both one-sided Kolmogorov-Smirnov statistics were computed, with Bonferroni correction applied.

The density of synapses at each path distance bin x was also quantified relative to the overall density of synapses, i.e.

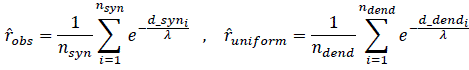

This was calculated for both the observed and above-mentioned resampled synapses, and a bootstrap-based Kwiatkowski-Phillips-Schmidt-Shin (KPSS) test for level stationarity was used to test if synapse density for each input region significantly varied with path distance, again using the recalculated KPSS statistic from the resamples as the null distribution.

The nearest neighbour distance (NND) for each synapse was quantified as the average path distance to the nearest distal neighbour and nearest proximal neighbour on the same neurite^99^. To test whether synapses tended to be clustered closer together than at random, a bootstrap-based Kolmogorov-Smirnov test was used to compare the distribution of NNDs to the distribution that would be expected if synapses were uniformly distributed across dendritic branches (while accounting for path distance-dependent variations in synapse density). This reference distribution could not be analytically derived and was therefore estimated using Monte Carlo methods as the mean distribution observed in 10000 resamples. Here, the locations of the *k* observed synapses for each cell were resampled by sampling *k* points with replacement from its dendritic tree, at a probability that was weighted by the synapse density observed at the point’s corresponding path distance bin, but uniform with respect to branch identity i.e. two points at the same path distance from the soma on different dendritic branches had equal probability of being sampled.

### Characterisation of biophysical properties of dPAG dendrites

#### Mice

For dual dendritic and somatic whole-cell recordings, male or female vGluT2-eYFP mice between 5-10 weeks of age were used. For holographic stimulation experiments, male or female vGluT2-Cre mice of 4-5 weeks of age were injected with a 1:1 mixture of AAV2-CAG-Flex-Flp and AAV-EF1a-fDIO-hChR2(H134R)-EYFP in dPAG, and given 2 weeks for viral expression before use in slice electrophysiology.

#### Preparation of acute coronal PAG slices

Slice preparation procedures and solution compositions were as previously described in^31^. Briefly, mice were intracardially perfused with ice-cold slicing ACSF under terminal isoflourane anaesthesia and decapitated. The brain was quickly extracted and sectioned into 250um thick coronal slices at the level of PAG using a vibratome (Leica VT1200S, Leica Biosystems) while immersed in ice-cold slicing ACSF. Slices were transferred to a recovery chamber containing slicing ACSF at near-physiological temperature (32°C) for 10 minutes, followed by transfer to a separate recovery chamber containing holding ACSF at room temperature for at least an hour before recordings. For recording, slices were transferred to a submersion type recording chamber (Scientifica) that was continuously perfused with room temperature recording ACSF using a peristaltic pump (PPS2, MultiChannel Systems). All solutions were constantly bubbled with 95% O2 and 5% CO2 and adjusted to a pH of 7.3-7.4 and osmolarity of 295-305 mOsm/kg.

Compositions of solutions were:

NMDG-based slicing ACSF (in mM): 96 NMDG-Cl, 2.5 KCl, 1.25 NaH2PO4, 25 NaHCO3, 20 HEPES, 25 D-glucose, 5 sodium L-ascorbate, 2 thiourea, 3 sodium pyruvate, 3 myo-inositol, 12 N-acetyl-L-cysteine, 0.01 taurine, 0.5 CaCl2, and 10 MgSO4.

Holding ACSF (in mM): 87 NaCl, 2.5 KCl, 1.25 NaH2PO4, 25 NaHCO3, 20 HEPES, 25 D-glucose, 5 sodium L-ascorbate, 2 thiourea, 3 sodium pyruvate, 3 myo-inositol, 12 N-acetyl-L-cysteine, 0.01 taurine, 2 CaCl2, and 2 MgSO4.

Recording ACSF (in mM): 125 NaCl, 2.5 KCl, 1.25 NaH2PO4, 26 NaHCO3, 5 HEPES, 10 D-glucose, 2 CaCl2, 1 MgSO4, 2 ascorbic acid, 2 sodium pyruvate, and 3 myo-inositol.

#### Dual dendritic and somatic whole-cell recordings

Recordings were made at the soma and dendrite of VGluT2+ dPAG neurons under visual guidance using infrared oblique illumination on an upright SliceScope Pro 1000 (Scientifica) equipped with a 60× water-immersion objective (LUMPlanFLN, NA1.0, 2 mm working distance, Olympus). VGluT2+ neurons were identified based on fluorescence from EYFP expression using 490 nm LED illumination (pE-100, CoolLED). Patch pipettes were pulled from standard-walled filament-containing borosilicate glass capillaries (GC150F-10, 1.5 mm outer diameter, 0.85 mm inner diameter, 100 mm length, Harvard Apparatus) using a horizontal micropipette puller (P1000, Sutter), to a resistance of ∼5 MΩ or ∼25 MΩ for somatic and dendritic recordings respectively. Pipettes were backfilled with intracellular solution adjusted to pH of 7.3-7.4 and osmolarity of 285-295mOsm/kg, containing (in mM): 130 KMeSO3, 10 KCl, 10 HEPES, 4 NaCl, 4 Mg-ATP, 0.5 Na2-GTP, 5 NaPhosphocreatine, 1 EGTA. 50 μM Alexa 594 was additional included to allow for visualisation of dendrites of the filled, recorded cell using 564 nm LED illumination. Recordings were made using EPC 800 amplifiers (HEKA), low-pass filtered at 5 kHz and digitised at 25 kHz using a PCIe-6353 board (National Instruments) controlled using custom software in LabVIEW.

To obtain dual somatic and dendritic recordings, the soma was held in gigaohm seal configuration without breaking into the cell while another pipette was used to target the dendrite of the same cell. This minimized occurrences of losing somatic access due to movements in the slice from the dendritic pipette. Once a gigaohm seal was obtained at the dendrite, brief suction pulses were applied first at the somatic pipette and then the dendritic pipette. Whole cell capacitance and series resistance were estimated and compensated for. Input resistance and series resistance were monitored continuously throughout the experiment using brief 250 ms, –40 pA hyperpolarising test pulses.

At either of the somatic and dendritic electrodes, current of the following waveforms were injected with an interstimulus interval of 1.5 s: 1) 3 repeats of 500 ms current steps, in 20 pA steps from –40 pA to 200 pA, 2) 40 repeats of 20 or 40 pA excitatory postsynaptic current (EPSC)-shaped currents, with an instantaneous rise and exponential decay at a time constant of 2 ms.

#### Holographic stimulation

Whole-cell recordings were made at the soma of ChR2-expressing, VGluT2+ dPAG neurons under visual guidance using infrared differential interference contrast illumination on an upright SliceScope Pro 6000 (Scientifica) equipped with a 40× water-immersion objective (LUMPlanFLN, 0.8 NA, 3.3 mm working distance, Olympus), following procedures described above. Once filled with Alexa 594 dye, dendrites were visualised with a Yokogawa CSU-X1 spinning disk confocal system equipped with 561 nm laser line (Vortran Stradus VersaLase) and emission filter centred at 617 nm. This was used to guide the placement of holographic stimulation spots over dendrites belonging to the recorded neuron. Holographic stimulation was achieved with a 1040nm laser (femtoTrain 1040-5, Spectra-Physics) coupled to a spatial light modulator (LCOS-SLM X10468, Hamamatsu; installed as Phasor system, 3i Intelligent Imaging Innovations) whose phase is programmatically controlled (Slidebook 6, 3i Intelligent Imaging Innovations) to generate precise spatial patterns at the sample. ChR2 was excited at a single diffraction-limited stimulation point with a 10ms pulse repeated 5 times at an interstimulus interval of 1 s, while voltage responses were recorded at the soma.

Stimulation location was confirmed with simultaneous confocal imaging of the Alexa594 dye excited by the holographic stimulation. The length of dendrite stimulated during each point stimulation was estimated from the full width at 1/e^2^ of the maximum excited Alexa594 fluorescence, corresponding to the effective excitation beam waist diameter, measured as 2.99 ± 0.155 μm. The full-width half-maximum of the excited fluorescence spot was 1.52 ± 0.0758 μm. The stage was then moved to re-position the holographic stimulation spot at a different dendritic location that could either be distal or proximal to the previous stimulation spot or on a different dendrite branch, in a randomised order. At the end of the recording, an image stack of the recorded and filled neuron was acquired to allow for reconstruction of the stimulation locations.

#### Analysis of voltage responses

Voltage responses were averaged across repetitions. Response amplitudes were quantified as the peak membrane voltage change for transient signals (action potentials, responses to EPSC shaped current injections and holographic stimulations), or the average steady-state response for step current injections. Attenuation was quantified as the ratio between the response at the passive recording electrode and the response at the current injection electrode, for both forwards and backwards propagation along the dendrite. The effective length constant (λ_eff_) was derived from the exponential fit of experimental attenuation values. Rise times were quantified as the time taken to rise to 1-1/e (approximately two-thirds) of the peak response. Half width of action potentials was quantified as the width of the spike at half of its peak value (full width at half maximum), and latency as the time between peaks in the somatic and dendritic recordings. Input resistance (R_i_) of a cell was quantified as the steady state voltage response to a hyperpolarising step current injection divided by the magnitude of the current injection, and the membrane time constant (τ_m_) as the time taken for the membrane voltage to drop to 1/e of the steady state response.

The path distance of the dendritic electrode site was estimated from 2D widefield epifluorescence snaps of the Alexa 594-filled cells in which the soma and dendritic pipette fell in the same field of view. To estimate the path distances of holographic stimulation sites, the 2D image snaps obtained during stimulation were combined with the corresponding stage coordinates and registered to the 3D image stack acquired at the end of the experiment using a modification of SHARP-Track. This positioned the stimulation sites in the 3D image stack, and dendrites of the recorded cell in the 3D image stack were then reconstructed.

### Biophysical modelling using NEURON

#### Fitting to experimental data

For dPAG simulations, reconstructions from the holographic stimulation experiments were used to generate multi-compartmental models in NEURON. Specific membrane capacitance was set to 1 μF/cm2 throughout the model, as spines were rarely observed. Specific membrane resistance (R_m_) and specific axial resistance (R_a_) were obtained by finding the combination of parameters that produced the best fit to the experimentally obtained input resistance and membrane time constant, experimentally observed forwards and backwards steady-state and transient attenuation values from the dual somatic and dendritic recordings, and experimentally observed rise times from the holographic stimulation experiments. Simulations of the experiments were done by replicating the various somatic and dendritic current injection waveforms or ChR2 photostimulation paradigms. ChR2 was simulated as a membrane density mechanism with dynamics based on a four-state Markov model that has two open and two closed states to capture the activation, inactivation and deactivation of ChR2 (obtained from^100^), at a uniform peak conductance density of 3 mS/cm^2^ across the cell, and 3 μm of dendrite photostimulated at 470nm at an uniform irradiance of 1.25 μW/μm^2^ for each dendritic point stimulation. Peak conductance density is difficult to estimate but typically modelled in the order of 1 mS/cm^2^; possible range based on expression density (1.3 x 10^10^ channels/cm^2^ for bacteriorhodopsin in oocytes^101^; estimated at 4.41 x 10^12^ channels/cm^2^ for ChR2 in neuronal membrane^102^) and single channel conductance (50-250 fS^103^) is 0.65 – 1100 mS/cm^2^.

Parameter grid search found that R_a_ values between 105 – 125 Ω cm and R_m_ values between 18400 – 18500 Ω cm^2^ produced membrane properties, effective length constants and rise time dependence that fell within the confidence intervals of all experimentally estimated values (R_i_, τ_m_, steady-state backwards λ_eff_, steady-state forwards λ_eff_, transient backwards λ_eff_, transient forwards λ_eff_, slope of rise time against path distance). R_a_ and R_m_ were thereafter set to 115 Ω cm and 18400 Ω cm^2^ respectively.

Dentate gyrus granule cell models (taken from^65^) were passive cells with an average R_m_ of 38000 ± 2300 Ω cm^2^ and R_a_ of 194 ± 24 Ω cm and no active conductances.

#### Simulations of synaptic integration

For synaptic integration simulations, synaptic conductance changes were simulated using the sum of two exponential functions with τ_rise_ = 0.2 ms and τ_decay_ = 2 ms, reversal potential of 0 mV, and peak conductances of 2 nS to produce ∼3mV somatic depolarisation when placed at the soma (in both dPAG and granule cell models).

To investigate the change in somatic response size with path distance of the synapse from the soma, single synapses were placed at each dendritic location and stimulated while simulated responses were recorded at the soma. Responses were normalized to the response generated in response to a synapse on the soma itself. Length constants were derived from the exponential fit of simulated normalised peak somatic response magnitude. Average expected response sizes 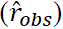 given synapses at path distances (*d*_*syn*) experimentally observed in anatomical reconstructions (from Chapter 4) were estimated for each reconstructed cell using the length constants (λ), and normalized to expected response sizes given a uniform synapse distribution 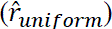, calculated respectively as:

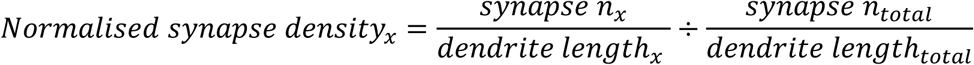

The normalized average expected response size 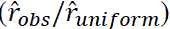 was calculated in a similar way for synapses resampled under the null hypothesis of uniform synapses 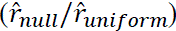, to assess significance of difference from the response size for uniform synapses. Values closer to 1 indicate greater similarity to the response size for uniform synapses, and the distance to 1 given PAG versus DGGC length constants were compared using the Wilcoxon signed-rank test. Normalized average expected response sizes were log-transformed for statistical tests, to ensure symmetrical distribution about log(1).

To investigate spatial summation, synapses were placed at all possible pairs of dendritic locations belonging to the same primary dendrite (i.e. sharing the same somatic branch origin), and the somatic response to stimulating both synapses simultaneously was normalized to the linear sum of somatic responses to individual stimulation of either synapse. Variability in spatial summation was quantified as the range (maximum-minimum) of the normalized response observed across all locations for each cell. Independent samples t-test was used to compare the variability in dPAG cells versus DG GCs.

To investigate temporal summation, a single synapse was placed at each dendritic location and stimulated twice with an interstimulus interval (ISI) ranging from 0 ms (i.e. simultaneous) to 100 ms. Responses were normalised to the response generated when the synapse was stimulated once. Variability in temporal summation was quantified at each ISI, as the range (maximum-minimum) of the normalized response observed across all locations for each cell. The temporal summation curve at each dendritic location was additionally described using the ISI that evoked peak normalised response as the peak ISI and the full-width half maximum as the width of temporal summation window. The variability in peak ISI and temporal summation window width were quantified as their respective ranges across all locations for each cell. Independent samples t-test was used to compare the variability of temporal summation at each ISI, peak ISI and window width in dPAG cells versus DG GCs.

### General statistical analysis

Unless otherwise specified, Bonferroni correction was applied in all cases involving multiple comparisons.

### Data and code sharing

Data and code will be shared at the time of publication and are available before that upon request.

## Author contributions

Y.L.T. performed the experiments; Y.L.T., A.T., N.Z. and D.C. analysed the data; T.B., Y.L.T. and D.C., designed the experiments; T.B. and Y.L.T. conceived the project. T.B., D.C. and Y.L.T. wrote the manuscript.

## Acknowledgments

This work was funded by a Wellcome Senior Research Fellowship (214352/Z/18/Z), by the Sainsbury Wellcome Centre Core Grant from the Gatsby Charitable Foundation and Wellcome (GAT3755 and 219627/Z/19/Z) and by a European Research Council grant (Consolidator #864912) (T.B.), Gatsby Unit/SWC Joint Research Fellowship in Neuroscience (D.C.), A*STAR National Science Scholarship (PhD; Y.L.T) and the SWC PhD Programme (Y.L.T.). We thank members of the Branco lab, in particular Y. Lefler, J. Bakermans, P. Iordanidou, D.S. Liu and S. Han, for help and discussions, and the SWC Advanced Microscopy facility, the SWC Neurobiological Research Facility, the SWC Viral Vector Core and the SWC FabLabs for technical support. We thank B. Cruz and G. Lopes for support with programming the data acquisition software.

