## Supplementary figures for "Presynaptic temporal dynamics flexibly set input weights in the mouse escape circuit"

**SUPPLEMENTARY INFORMATION**

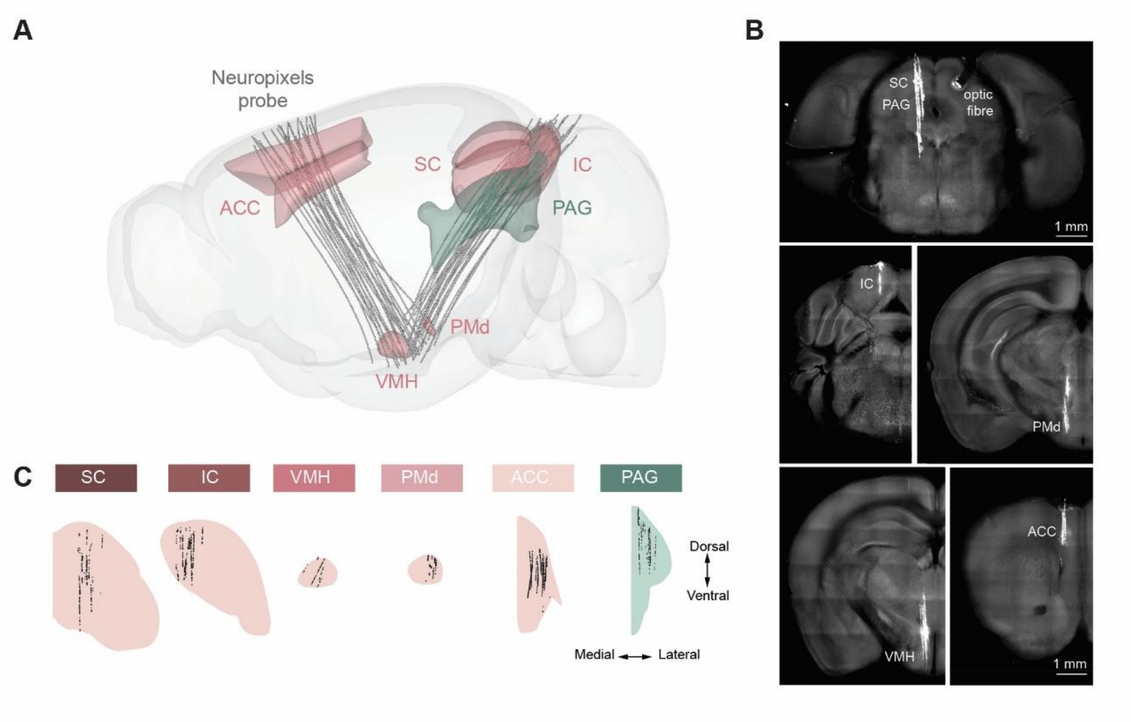


**Figure S1 - Location of targeted regions and recorded neurons across the brain**

**(A)** Reconstructed probe trajectories from all 6 mice. **(B)** Serial two-photon tomography images of an example mouse brain, with probe shanks in white. **(C)** Coronal view of reconstructed locations of recorded single units across all recording sessions, in each region.

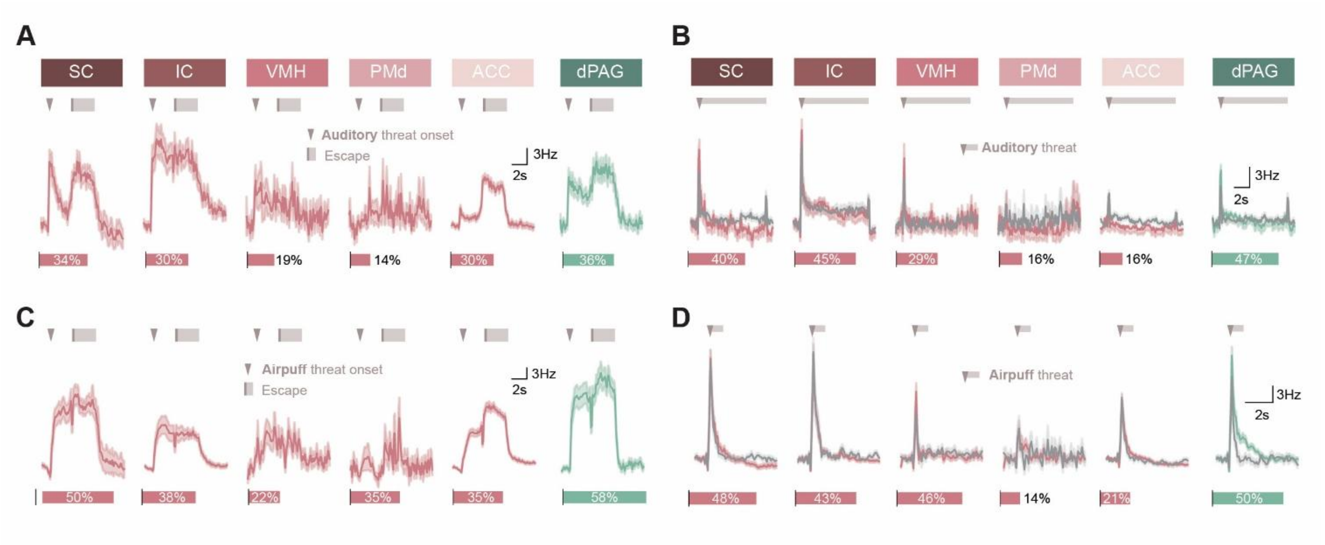


**Figure S2 - Escape and threat neural activity during auditory threat and air puff trials**

**(A)** Proportion of neurons active during escape from auditory threats (bottom bars), and average peri-escape PSTHs of the escape-excited cells (top), for each region. **(B)** Proportion of neurons activated by auditory threat presentation (bottom bars), and average peri-threat PSTHs of sound-excited cells (top), on escape (pink) and failure trials (grey). **(C-D)** Same as A and B respectively for air puff trials.

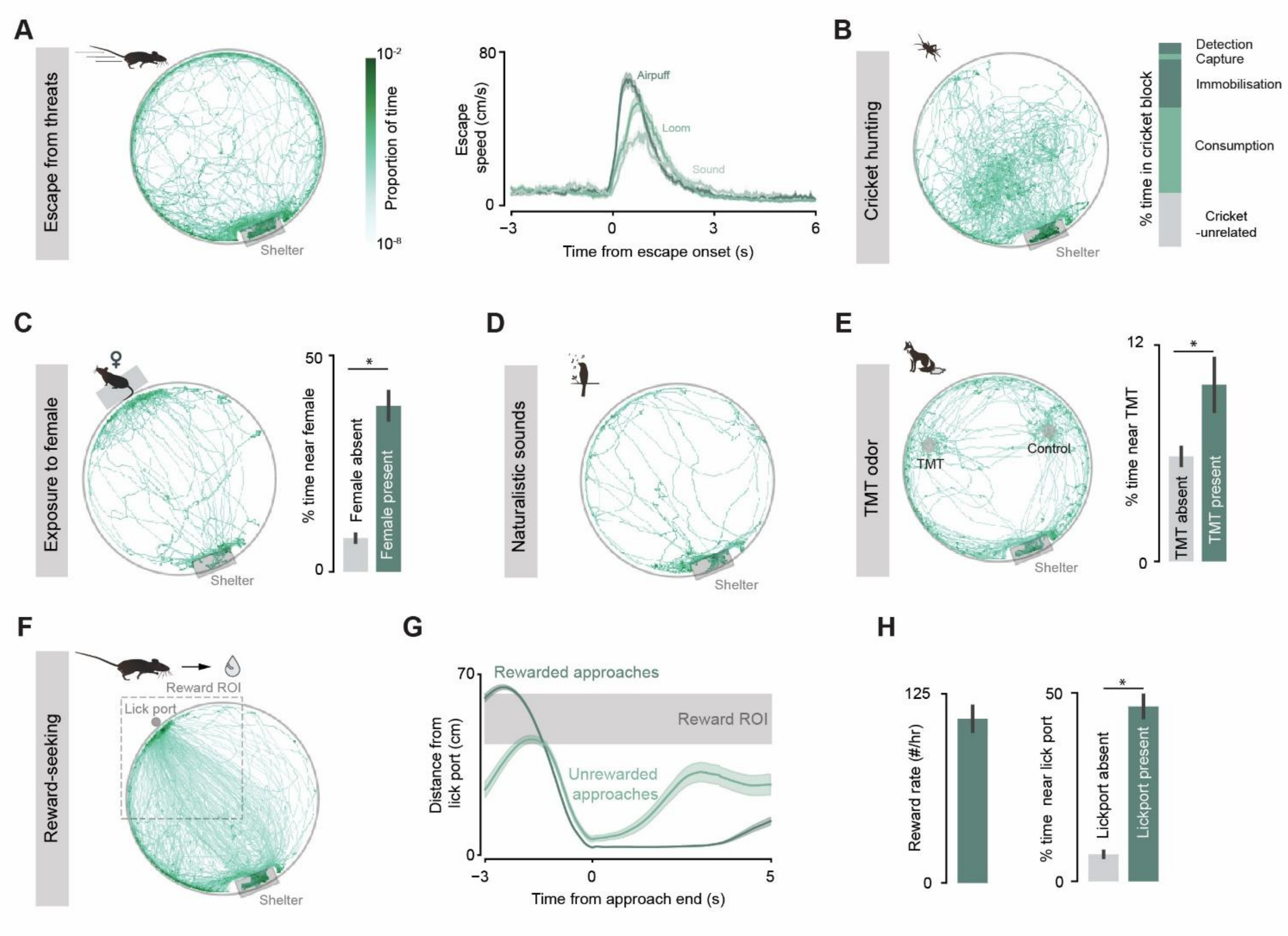


**Figure S3 - Behavioural quantification across experimental paradigms**

**(A)** Example heatmap of mouse location during escape block (left). Average speed traces aligned to escape onset (right) showing rapid run back to shelter upon threat. **(B)** Example heatmap of mouse location during cricket block (left) and session-averaged percentage of time spent on hunting-related behaviours during the block (right). Most time was spent on the different stages of the hunting sequence (prey detection: 5.5 ± 1.1 %, prey capture: 2.6 ± 0.7 %, attack: 23.7 ± 2.1 %, consumption: 41.8 ± 3.3 %); cricket-unrelated behaviours (26.4 ± 3.8 %). **(C)** Example heatmap of mouse location during female block (left), with female mouse positioned adjacent to the arena in an enclosed box with ventilation holes. Average percentage of time spent near the female mouse during the female block (38.4 ± 3.28 %), compared to similar locations during the escape block when the female was absent (7.79 ± 0.931 %; paired t-test, $p = 1.84 \times 10^{-9}$). **(D)** Example mouse location during sound block, during presentation of naturalistic sounds. **(E)** Example heatmap of mouse location during TMT block (left), with TMT-loaded filter paper placed in one of two petri dishes. Percentage of time spent near the TMT-loaded filter paper (right) during the TMT block (green; 9.66 ± 1.31 %), compared to similar locations during the escape block when TMT was absent (grey; 5.59 ± 0.460 %, paired t-test, p = 0.00535). **(F)** Example heatmap of mouse location during reward block (left). Lick port that dispenses condensed milk is added to arena, and reward delivery is contingent on approaches beginning from outside the reward ROI. Note loops formed as mouse shuttles between the lick port and shelter. **(G)** Average distance from lick port for rewarded approach trials (dark green) and unrewarded trials (light green), where mice initiate approach too close to the lick port and leave the lick port soon after arrival because no reward is delivered. **(H)** Session-averaged rate of rewards obtained (left; 108.4 ± 8.3 rewards/ hr) and time spent near the lick port location (right) during the reward block (green; 44.5 ± 2.83 %), compared to similar locations during the escape block when rewards were absent (grey; 6.57 ± 0.786 %; paired t-test, $p = 1.95 \times 10^{-13}$), showing engagement in reward-seeking.

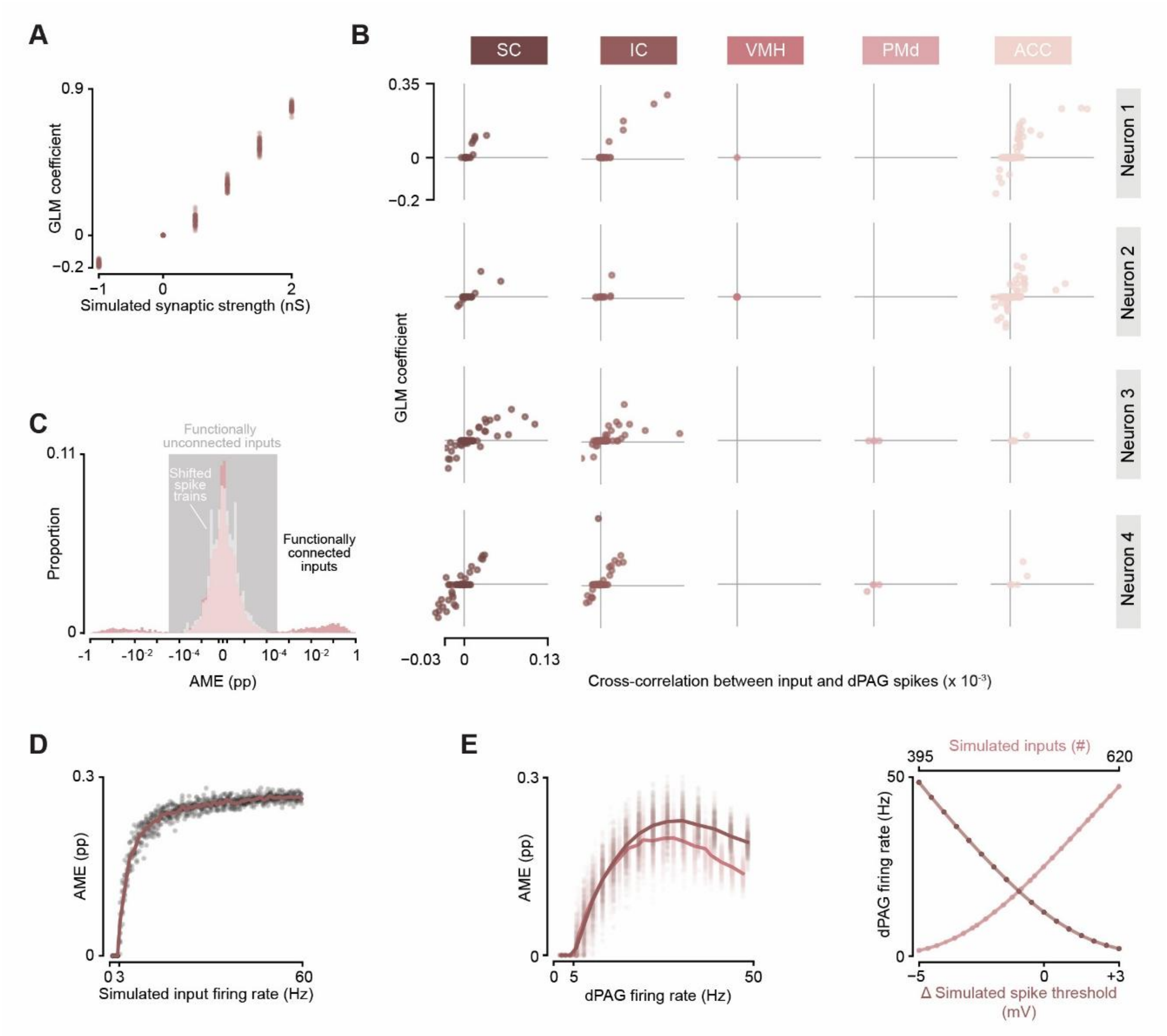


**Figure S4 - Additional GLM analyses**

**(A)** GLM coefficients for simulated inputs of increasing synaptic strength. **(B)** GLM coefficients for the inputs onto the four example dPAG cells in Figure 1I. **(C)** Histogram of average marginal effects (AMEs) across all inputs onto all well-predicted dPAG cells. AMEs are approximately symmetrically distributed around zero and trimodal, with a sharp peak near zero and broader peaks at larger positive and negative values. The near-zero peak (within $\pm 3 \times 10^{-4}$ pp) matches the distribution obtained when input spike trains are temporally shifted to eliminate any relationship with dPAG spiking; 98% of AMEs fall within this range and therefore correspond primarily to functionally unconnected pairs. **(D)** AMEs for simulated inputs at increasing firing rates. **(E)** AMEs for simulated inputs onto dPAG neurons of different firing rates (left). dPAG firing rate was varied by changing either the postsynaptic spike threshold (dark pink, right) or the number of convergent inputs driving the dPAG neuron (light pink, right).

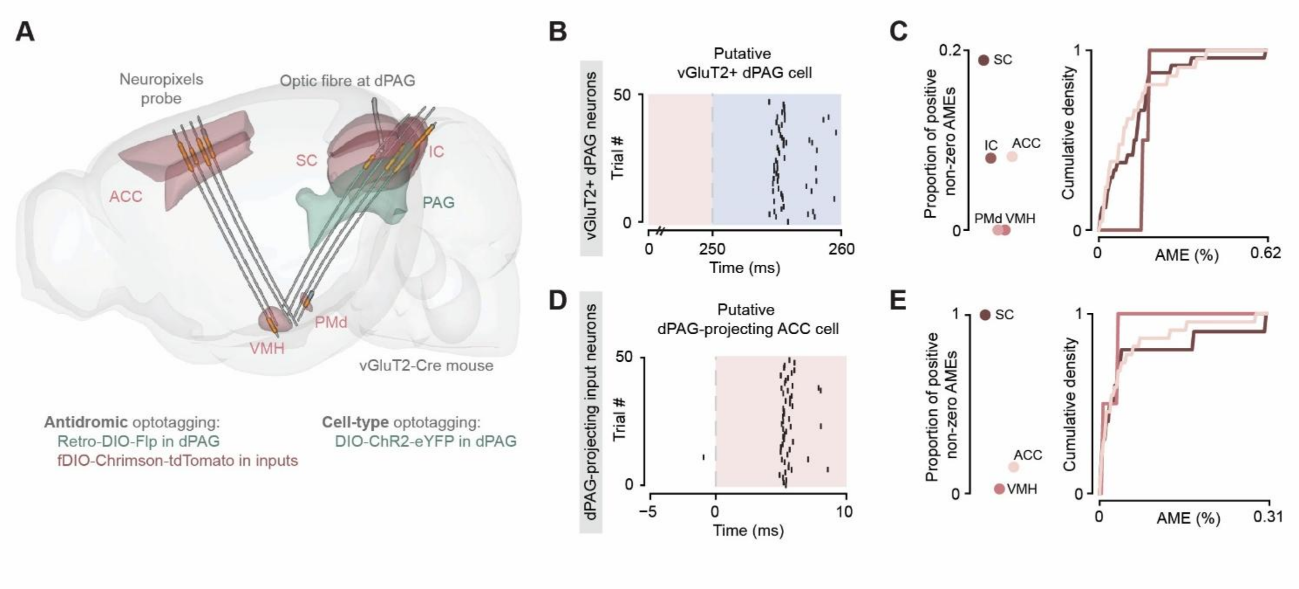


**Figure S5 - Optotagging of dPAG excitatory neurons and dPAG-projecting excitatory input cells**

**(A)** Schematic for viral and targeting strategy for recording from vGluT2+ dPAG cells and dPAG-projecting vGluT2+ or CaMKII+ inputs in SC, IC, VMH, PMd and ACC. Light was delivered at the dPAG via an optic fibre. **(B)** Example opto-tagged, putative vGluT2+ dPAG neuron with reliable light-evoked spikes at short latency and low jitter, following a 10 ms blue light pulse after 250 ms long red light pulse to trigger the depolarization block of surrounding Chrimson-expressing axons. **(C)** Proportion (left) and magnitudes (right) of positive non-zero AMEs of inputs to well-predicted putative vGluT2+ dPAG cells. **(D)** Example opto-tagged, putative dPAG-projecting ACC neuron with light-evoked spikes following a 10 ms red light pulse. **(E)** Proportion (left) and magnitudes (right) of positive non-zero AMEs of optotagged, putative dPAG-projecting inputs in SC, ACC and VMH.

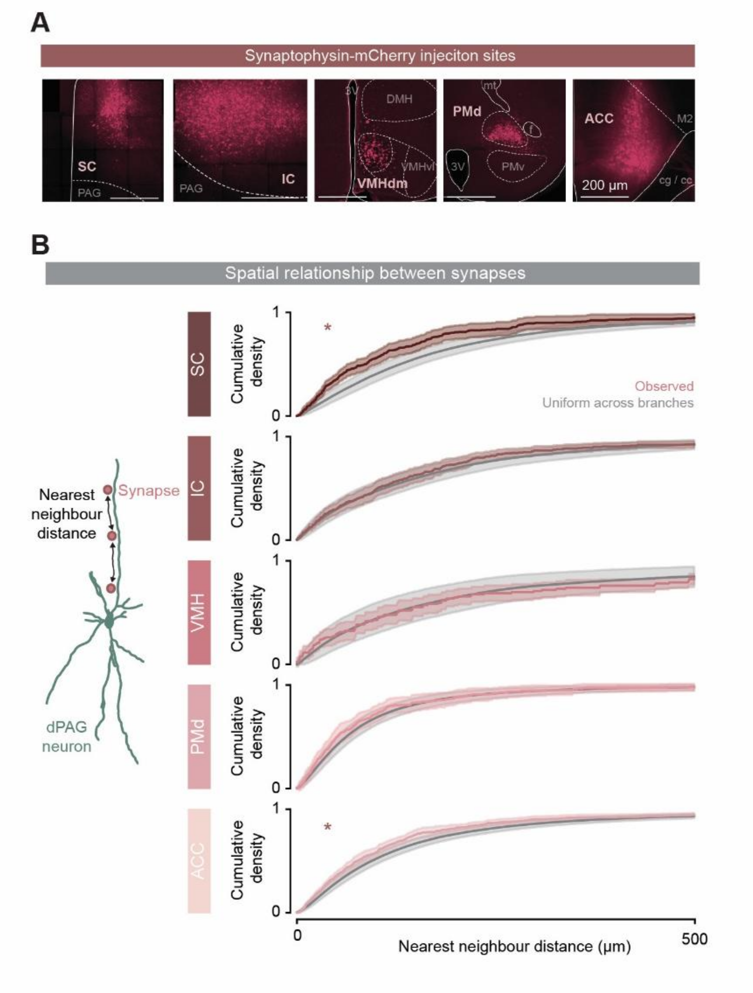


**Figure S6 - Additional quantifications of synapse organisation**

**(A)** Injection sites of synaptophysin-mCherry at SC, IC, VMH, PMd or ACC. (3V - 3rd ventricle; DMH - dorsomedial hypothalamic nucleus; f - fornix, mt - mammillothalamic tract; Cg – cingulum; cc - corpus callosum). **(B)** Schematic of nearest neighbour distance as a metric for synapse clustering (left), quantified as the average path distance to the nearest distal neighbour and nearest proximal neighbour on the same neurite. Empirical cumulative density functions (right) of observed nearest neighbour distance between synapses, pooled across cells (pink). Pink shading indicates weighted SEM, with each cell weighted by number of synapse pairs. Grey line shows expected nearest neighbour distances for synapses uniformly distributed across branches (and following observed path distance-dependent variations in synapse density), estimated using Monte Carlo methods as the mean of resamples (grey). Grey shading indicates 1-99 percentiles of resamples. Asterisks indicate significant difference from expected for uniform distribution across branches.

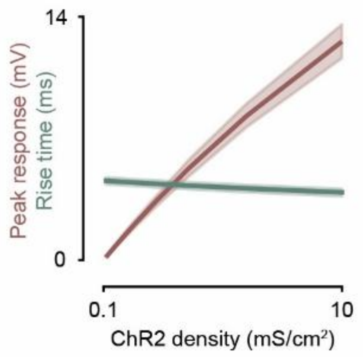


**Figure S7 - Invariance of response rise times with ChR2 expression**

Local peak ChR2-evoked response size (red) scales almost linearly with ChR2 conductance density in simulated dPAG cells, except for a mild saturation of driving force producing a slightly sublinear increase in photocurrents only at very high expression levels. In comparison, rise times (green) remain much more stable, with only a slight shortening of rise times resulting from the increased opsin conductance reducing membrane resistance and shortening the effective membrane time constant. A 100-fold increase in ChR2 conductance density from 0.1 mS/cm2 to 10 mS/cm2 results in a 76.2-fold increase in peak voltage amplitude, but only a 1.17-fold reduction in rise time.

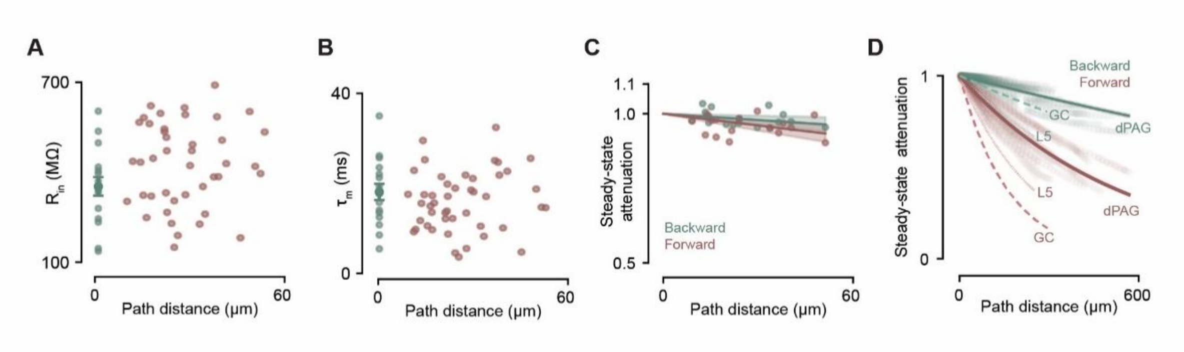


**Figure S8 - Additional quantifications of passive properties**

**(A-B)** Input resistance (A) and effective membrane time constant (B) at soma and dendrite, estimated from hyperpolarising step current injections. **(C)** Backward (green) and forward (pink) attenuation of steady-state responses. Attenuation is quantified as the ratio of voltage at the recording site to voltage response at stimulation site. Effective path lengths in the backwards and forwards directions were 1403 ± 446 µm (n = 15) and 747 ± 185 µm (n = 18) respectively. **(D)** Backward (green) and forward (pink) attenuation of steady-state responses in dPAG compartmental models, showing shallower decrease with path distance when compared to dentate gyrus granule cells and basal dendrites of layer 5 pyramidal neurons.

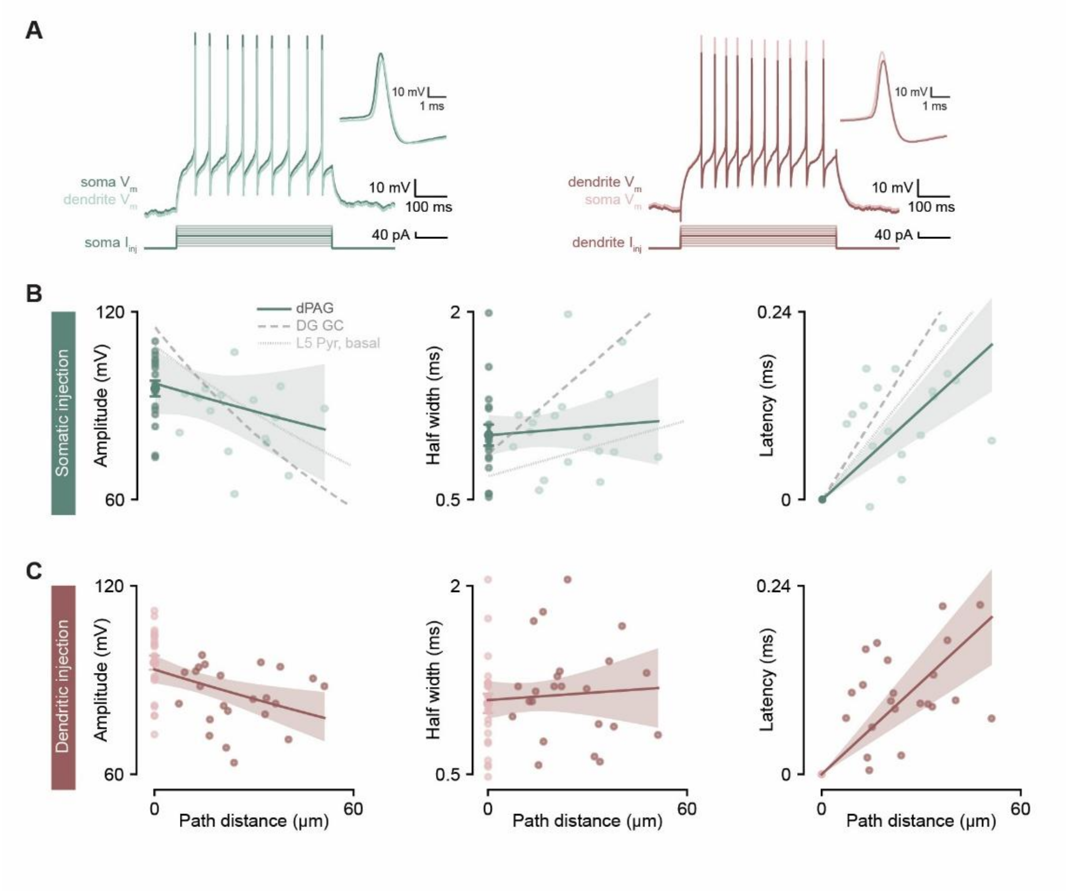


**Figure S9 - Action potential back-propagation in dPAG dendrites *in vitro***

**(A)** Somatic and dendritic voltage responses to depolarising step current injections at the soma (left, green) or dendrite (right, pink).

**(B-C)** Amplitudes (left), half widths (middle) and peak latencies (right) of back-propagating action potentials (bAPs) evoked by somatic (B, n = 18) or dendritic (C, n = 23) current injection. bAPs attenuate with a length constant of 313 ± 161 μm, broaden at a rate of 2.18 ± 4.68 μm, and are conducted at a velocity of 259 ± 46.6 μm.

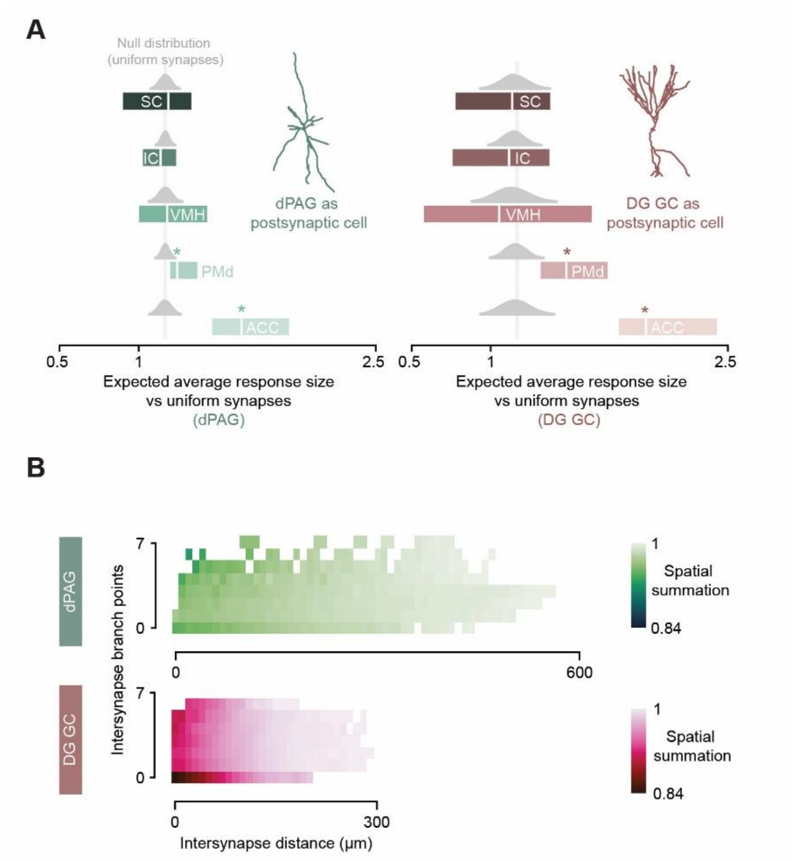


### Figure S10 - Synaptic integration of region-specific inputs in dPAG and dentate gyrus granule cell models

**(A)** Expected average response sizes to SC, IC, VMH, PMd and ACC inputs per postsynaptic cell, given the anatomically observed distributions of synapse path distances and either dPAG (left, green) or DG GC (right, pink) somatic response decay length constants, normalized to expected average response sizes for uniformly distributed synapses. Boosting of PMd and ACC inputs by proximal enrichment is significantly smaller with the dPAG length constant than the DG GC length constant (Wilcoxon signed-rank test, n = 17, p = $1.53 \times 10^{-5}$ for PMd inputs and n=9, p = $2.73 \times 10^{-2}$ for ACC inputs). **(B)** Heatmap of spatial summation in simulated dPAG cells (top, green) and dentate gyrus granule cells (bottom, pink), at varying number of branch points (y axis) and path distances (x axis) between synapse pairs.

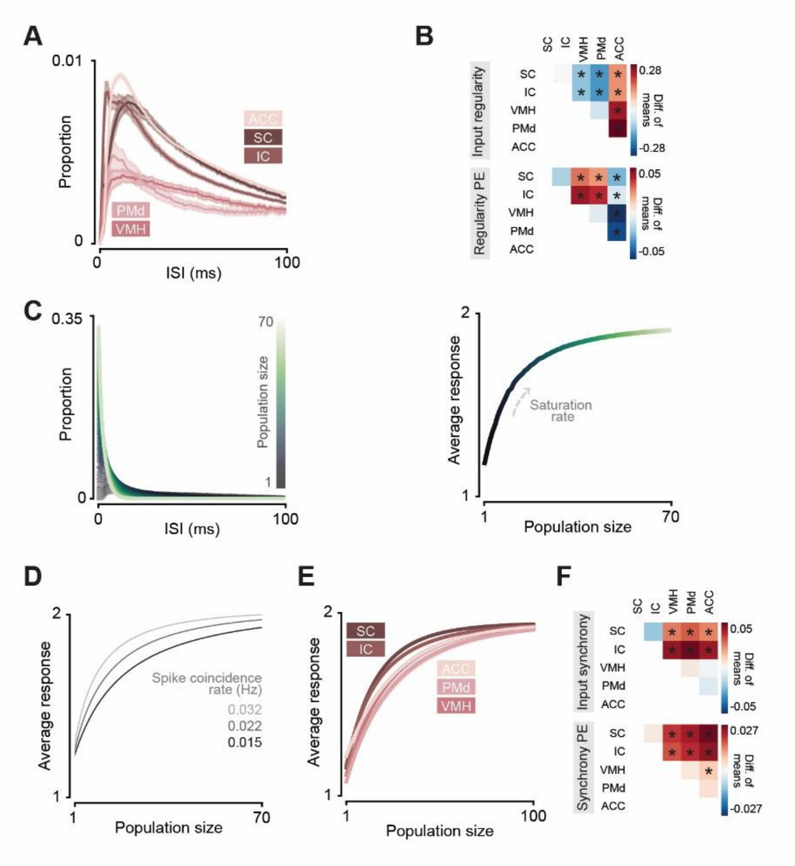


**Figure S11 - Additional analyses of presynaptic temporal statistics and their postsynaptic effects**

**(A)** Distributions of inter-spike intervals (ISI) within individual neurons, for each region. **(B)** Pairwise differences in mean gamma shape parameter α (top) and Poisson-normalised mean expected response size (bottom) between brain regions (row − column). Asterisks: significant differences (wild bootstrap test). **(C)** Estimating the postsynaptic effect of synchrony for an example cell. Left: the proportion of the cell's spikes occurring within 100 ms of another spike, considering either the cell alone (population size = 1, dark green) or together with increasing numbers of other simultaneously recorded cells from the same region (lighter green). Right: convolving each resulting ISI distribution with the temporal-summation curve yields the mean expected postsynaptic response at each population size. The average expected response saturates as all spikes of the cell's spikes fall within 1 ms of another spike in the population, at a rate depending on synchrony with the population. **(D)** Mean expected response as a function of population size, for simulated spike trains with increasing rates of coincident spikes between neurons (lighter grey indicates higher synchrony), showing faster saturation at higher synchrony. **(E)** Region-averaged curves of mean expected response as a function of population size. **(F)** Pairwise differences in mean cross-covariance at zero lag (top) and mean saturation rate constant (bottom) between brain regions. Asterisks: significant differences (wild bootstrap test).

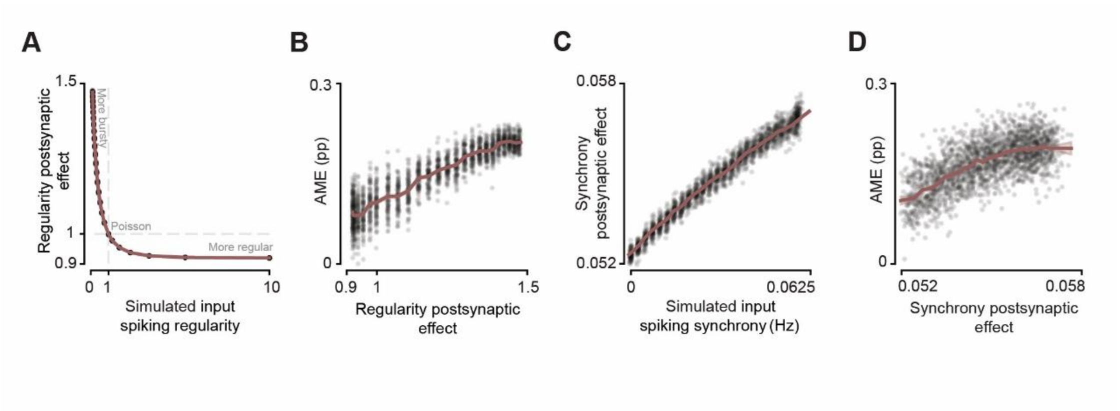


**Figure S12 - Presynaptic temporal statistics influence functional connectivity strength in simulated feed-forward networks**

**(A-B)** Poisson-normalized average expected responses (A) and GLM AMEs (B) assigned to simulated inputs with spike regularity varied by sampling ISIs from gamma distributions of varying shape parameter α. **(C-D)** Saturation rate constants of average expected response with population size (C) and GLM AMEs (D) assigned to simulated inputs with varying zero-lag inter-neuron synchrony.

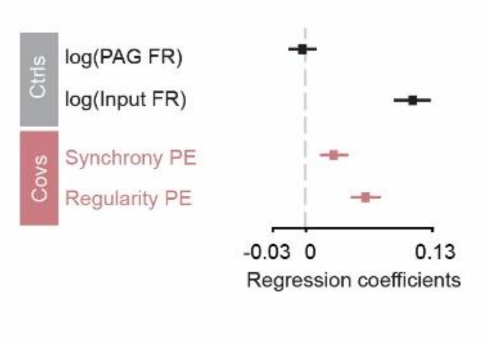


**Figure S13 – Quantification of GLM AMEs and temporal covariates during threat-only and motivational conflict block**

Regression coefficients of the fixed effects in a linear mixed model of AME magnitude with ACC-dPAG pairs as random intercepts (in 10 minutes bins). Fixed effects include time-varying dPAG and input firing rates as control variables (grey) and regularity and time-varying synchrony postsynaptic effect as temporal statistics covariates (pink). Error bars are bootstrap-based SEM.
